# Individualised electrophysiological neural field models for the assessment of thalamocortical function in disorders of consciousness: a multicentre study

**DOI:** 10.1101/2025.10.24.683919

**Authors:** Lín Cóngyŭ, Prejaas Tewarie, Naji L.N. Alnagger, Iván Mindlin, Glenn J.M. Van der Lande, Laouen Belloli, Steven Laureys, Lionel Naccache, Benjamin Rohaut, Aurore Thibaut, Jitka Annen, Olivia Gosseries, Pablo Núñez, Jacobo D. Sitt

## Abstract

Understanding the neural mechanisms underlying disorders of consciousness (DoC) remains a major challenge, particularly in distinguishing limited awareness in minimally conscious state (MCS) and complete unawareness in unresponsive wakefulness syndrome, also coined vegetative state (UWS/VS). In this multicentre study, we fitted a biophysically informed corticothalamic neural field model to high-density EEG data from two large independent datasets, comprising 203 UWS patients, 270 MCS patients and 74 healthy controls. We then used the fitted parameters to simulate EEG time series on a per-subject basis and compared empirical and simulated complexity metrics. The model reliably captured the spectral features across different states of consciousness and revealed reduced corticothalamic integrity in DoC patients that was more pronounced in UWS than in MCS, supporting the mesocircuit hypothesis. Furthermore, the simulated EEG reproduced the complexity patterns of the empirical recordings, with permutation entropy emerging as a sensitive marker capable of distinguishing between MCS and UWS for both real and simulated time series.

## Introduction

Disorders of consciousness (DoC) are a variety of clinical conditions in which patients who suffer from severe acquired brain injury experience impaired wakefulness and/or awareness ^1^. Among them, coma refers to a state of complete absence of both awareness and wakefulness ^2^. Usually within a month after injury, it can evolve into unresponsive wakefulness syndrome/vegetative state (UWS/VS), in which patients display signs of wakefulness such as eye-opening and reflexive movements, and/or the minimally conscious state (MCS), where patients additionally show minimal signs of awareness such as command following or visual pursuit ^3,4^. The emergence from the MCS (eMCS), encompasses those individuals who regain functional communication or purposeful objects use, and they are no longer considered to be DoC ^3^.

The thalamus is a subcortical structure that serves as a central communication hub between the subcortical regions, the cerebellum, and the cortex. It not only acts as a first-order relay pathway between the sensory input and the primary sensory cortex, but also has higher-order nuclei, that facilitate corticocortical interactions, possibly contributing to the modulation of conscious states ^5,6^. It has been suggested that the thalamus plays a crucial role in supporting conscious experience due to its ability to compress high-dimensional cortical activity patterns and then relay them back to the cortex as a lower-dimensional output. This efficient information compression mechanism is thought to support the unified, low-dimensional “essence” of conscious contents, *i.e.* how they are experienced as an integrated whole instead of disconnected features ^5^. Notably, the central thalamus is critically involved in many cases of DoC, as shown by the fact that direct injury to this structure can by itself produce disturbances in consciousness ^6,7^. Specific subnuclei are known to result in DoC after bilateral injuries ^8^, and corticothalamic integrity in acute DoC patients is associated with more favourable clinical outcomes ^9^. These findings are consistent with the mesocircuit hypothesis, which posits that after severe brain injury, a widespread deafferentation of the striatum and thalamus leads to a breakdown in communication between the cortex, basal ganglia, and central thalamus ^8^. Disruptions in the thalamocortical loops are common in many types of brain injury and are associated with a reduced ability to sustain cortical activity, impairing both arousal (via brainstem/basal forebrain) and awareness (via higher-order cortical areas) ^5,7,8^. Indeed, interventions such as deep brain stimulation (DBS) of the central thalamus have been shown to be effective in restoring both arousal and awareness in anesthetized non-human primates ^10^, and to increase cognitively mediated behaviours in MCS patients after traumatic brain injury (TBI) ^11–13^. Notable progress has also been made in thalamic low-intensity focused ultrasound (LIFU) intervention ^14,15^. Case reports describe patients demonstrating clinically significant increases in behavioural responsiveness following each treatment, while maintaining stable blood pressure, heart rate, and blood oxygen saturation. Additionally, pharmacological interventions such as the dopaminergic agent amantadine were also associated with recovery in UWS and MCS patients ^16^. According to the mesocircuit hypothesis, amantadine may be effective by promoting dopaminergic input to the striatum, and thus enhancing thalamocortical synaptic activity ^8^. Thus, understanding patient-specific impairments in thalamocortical circuits could be key to enabling targeted interventions on DoC patients aiming at restoring arousal and awareness.

Assessing consciousness is inherently challenging because it is not a unitary construct and lacks a universally agreed-upon definition. In scientific investigations, however, it is often operationalized via two dimensions: arousal (the level of wakefulness) and awareness (the content of experience). These dimensions provide a practical framework for evaluating and classifying states of consciousness and related disorders ^17^. Moreover, since subjective experience cannot be directly measured, assessment of the states of consciousness relies on observable features. These include behaviour, assessed via standardized scales such as Coma Recovery Scale-Revised (CRS-R) ^18^, as well as neuroimaging evidence derived from techniques like magnetic resonance imaging (MRI), positron emission tomography (PET), and electroencephalography (EEG) ^19^. EEG stands as a powerful tool for the study of consciousness, given its high temporal resolution, non-invasiveness, and feature-rich spectral content ^20–22^. Its diagnostic significance is highlighted by the European Academy of Neurology’s recommendation to incorporate quantitative analysis of high-density EEG into multimodal assessments to distinguish between UWS and MCS ^19,23^.

While obtaining direct measures of thalamocortical activity is not possible with non-invasive EEG, models based on neural field theory (NFT), which model the non-linear dynamics of a large population of neurons ^24^, have been successfully used to predict the power spectrum of EEG. In particular, Robinson and colleagues ^25–27^ developed a physiologically informed NFT model that has repeatedly demonstrated its ability to accurately replicate empirically observable phenomena through corticothalamic populations and their interactions. Among others, they have modelled sleep and wake states ^28^, evoked response potentials ^29^, sleep spindles ^30^, seizures ^31^, and age-related EEG changes ^32^. Furthermore, the model parameters can be fitted to empirical spectra to determine biophysical properties of the thalamocortical loops, as they relate directly to the physiological system. Previous research has attempted to use the model to distinguish between conscious and unconscious states through the model parameters after fitting to the subject’s power spectrum, including wake and REM states, as well as UWS, MCS and eMCS ^33^. While they were successful in differentiating between conscious and unconscious states, namely wakefulness versus REM versus N3 sleep, and healthy participants versus patients with DoC, the parameter space could not distinguish between brain injury subgroups.

Numerous studies have shown that the complexity of brain signals is tightly linked with conscious experience, indicating that for a system to be able to support consciousness, it must be able to both integrate and differentiate information efficiently ^34^. Therefore, to validate the capacity of the corticothalamic model to reconstruct the neural dynamics underlying consciousness, we proposed to test its outputs against established neural complexity measures. Lempel-Ziv complexity (LZC), an algorithm that determines the compressibility of a signal by measuring diversity in signal activity patterns, has been successfully used on spontaneous EEG to show a sharp decline in differentiation during anaesthesia 40in humans ^35^ and in animal EEG and electrocorticography (eCoG) ^36^. Moreover, by measuring the LZC of the response of the cortex to transcranial magnetic stimulation (TMS), researchers have used the perturbational complexity index (PCI) to distinguish between conscious and unconscious states reliably ^37^. Additionally, permutation entropy (PE), a measure of signal complexity that quantifies signal irregularity and complexity by calculating the entropy of the frequency of each possible permutation of symbolic sequences for a specific embedding dimension ^38^, and spectral entropy (SE), a measure of the predictability or uncertainty of the power spectrum distribution ^39,40^, have been successfully used to differentiate between UWS and MCS patients, further supporting EEG complexity as a key index of consciousness ^41^. Based on this body of evidence, we hypothesized that a valid model should produce simulations mirroring these empirical findings.

In the present study, we aim to leverage two large independent high-density EEG datasets of healthy controls and patients with DoC (namely UWS and MCS) to determine the features of the model’s parameter space that would reliably distinguish between DoC subgroups, *i.e.* MCS and UWS, in addition to differentiating between healthy controls and DoC patients. We also investigated how the model’s performance is influenced by inter-patient variability, such as CRS-R index ^42^ and time since onset (TSO). Additionally, we examined the frontal and parietal cortex in addition to the averaged whole brain activity, as the Global Neuronal Workspace (GNW) hypothesis suggests a fronto-parietal network plays a crucial role in making information accessible for conscious experience ^43,44^. In contrast, the Integrated Information Theory (IIT) suggests that the content of consciousness is maintained by neural activity within a temporo-parietal-occipital “hot zone”, while the prefrontal cortex is more involved in conscious access ^45,46^. This ongoing debate highlights the need for further investigation into the distinct functional contributions of different brain regions to consciousness ^45,47–50^. Furthermore, we evaluated the model’s ability to reproduce not only the spectral features of the empirical EEG, but also their complexity profiles, demonstrating the model’s robustness in capturing multiple dimensions of the data.

The key objectives we addressed were the following: (i) we successfully fitted a corticothalamic neural field model to the EEG power spectrums of UWS, MCS, and healthy controls from two independent datasets; (ii) the model revealed distinct patterns of impaired corticothalamic circuits in patients with DoC, highlighting difference between MCS and UWS, in addition to comparing brain activity between patients and healthy controls; and (iii) we evaluated the model’s ability to simulate and recreate the brain complexity of the empirical recordings based on the biophysically-informed fitting of the parameters. These results demonstrate that the model captures both frequency-domain and dynamic characteristics of brain activity, providing a valuable framework for interpreting the underlying neurophysiological mechanisms of consciousness

## Materials and Methods

### Subjects

This retrospective study involved two independent datasets collected from the University and University Hospital of Liège, as well as the Pitié-Salpêtrière Hospital in Paris. Ethical approval for the study was granted by the Ethics Committees of both institutions, in accordance with the Declaration of Helsinki. Written informed consent was obtained from the legal representatives of all patients and from the healthy control participants. Clinical evaluations were conducted using the CRS-R, with the final diagnosis determined by the highest CRS-R score obtained across multiple assessments (a minimum of five in the Liège dataset and a minimum of three in the Paris dataset^51^). All assessments were administered on separate days by experienced clinicians.

EEG recordings in the Liège dataset were previously collected while participants were at rest ^52^. In contrast, the Paris dataset included task-related EEG data obtained using the auditory “Local-Global” (LG) paradigm ^53^, designed to explore both conscious and unconscious auditory processing. This paradigm presents sequences of five tones, with the final tone either matching the previous ones in pitch (local standard) or differing from them (local deviant). Additionally, a second level of regularity is introduced based on the overall frequency of the sequences, with 80% classified as global standards and 20% as global deviants.

We removed 1 MCS subject from the Liège dataset and 3 UWS subjects from the Paris dataset due to excessive signal noise. Eventually, the Liège dataset included 37 healthy controls and 120 patients with prolonged DoC (>28 days post-injury), comprising 33 UWS patients and 86 MCS patients. Portions of this dataset have been used in previous research ^21,33,52,54^. The Paris dataset consisted of 354 prolonged DoC patients: 170 UWS, 184 MCS, and 37 healthy controls. This dataset has also been previously used ^22,41,55–57^. The demographic details of both datasets are summarized in Tables 1 and 2.

**Table 1.** Socio-demographic and clinical data for the Liège dataset. Statistically significant between-group differences are marked with asterisks ∗p < 0.05 (Mann-Whitney *U*-test for age, chi-squared test for sex). m: mean; std: standard deviation; M: male; F: female; N/A: not applicable. CRS-R: Coma Recovery Scale – Revised.

| Data | Group |  |  |
| --- | --- | --- | --- |
|  | UWS | MCS | Controls |
| Number of subjects | 33 | 86 | 37 |
| Age (years) (m±std) | 42.25±15.04 | 40.95±17.20 | 43.50±15.63 |
| Sex (M:F:Missing) | 19:14:0 | 52:30:4 | 20:15:2 |
| CRS-R total score (m±std) | 6.39±1.34 | 11.58±4.81 | N/A |
| Anoxia (%) | 57.57 | 22.98 | N/A |
| Stroke (%) | 3.03 | 3.44 | N/A |
| Traumatic brain injury (%) | 24.24 | 45.97 | N/A |
| Other (%) | 15.15 | 18.60 | N/A |
| Time since onset (months) (m±std)* | 28.87±36.55 | 40.25±44.16 | N/A |

**Table 2.**
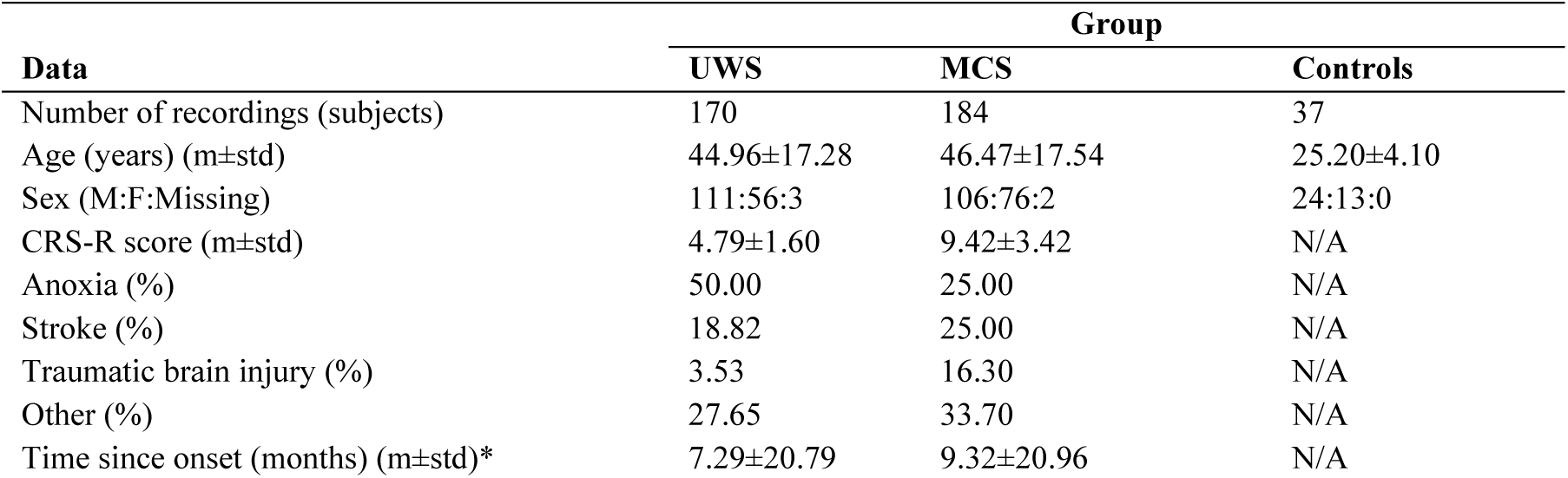
Socio-demographic and clinical data for the Paris dataset. Statistically significant between-group differences are marked with asterisks ∗p < 0.05 (Mann-Whitney *U*-test for age, chi-squared test for sex). m: mean; std: standard deviation; M: male; F: female; N/A: not applicable. CRS-R: Coma Recovery Scale – Revised.

### Electroencephalographic recordings

EEG recordings for both datasets were obtained using a 256-channel high-density EEG system (EGI, Electrical Geodesics Inc.) with a sampling rate of 250 Hz. During the recording sessions, subjects remained awake with their eyes open. Face and neck electrodes were removed, keeping a total of 183 channels.

In the Liège dataset, resting-state EEG was acquired for 20-30 minutes. The preprocessing pipeline included multiple steps. Initially, a 6th-order Butterworth high-pass filter at 0.5 Hz and an 8th-order IIR Butterworth low-pass filter at 100 Hz were applied, with notch filters at 50 and 100 Hz to remove power line interference. The EEG signal was then segmented into 2.2040-second epochs. Bad channels were identified through visual inspection of the raw time series, while noisy segments were detected by transforming the data into 20 principal components (PCA) and visually inspecting them. Independent component analysis (ICA) was performed to remove artifacts related to non-neural activity. Spherical interpolation was used to reconstruct rejected channels, and the final dataset was re-referenced to the average reference.

In the Paris dataset, EEG data were segmented into epochs ranging from −200 ms to 1336 ms relative to the onset of the first sound. Preprocessing was entirely automated. A band-pass filter with a bandwidth of 0.5 to 45 Hz was applied using a 6th- and 8th-order FFT-based Butterworth filter. Epoch lengths were set to 1.5 seconds. Adaptive outlier detection was then used to identify and exclude noisy epochs ^58^. Finally, the data were re-referenced using an average reference, and baseline correction was performed. Although task-based and resting-state EEG capture evoked and spontaneous neural activity respectively, studies demonstrate that consciousness markers derived from task-based paradigms can generalize to the resting state ^20,22^. This validates our approach of treating the LG paradigm data as a pseudo-resting state condition, incorporating all epochs of the recording regardless of the sequence.

### Neural field model

In a neural field model, the neural population activity is simulated by the mean firing rate of an afferent population within a node corresponding to a brain region. The model used in this study includes 4 interconnected populations within each node: cortical excitatory (*e*), cortical inhibitory (*i*), thalamic relay (*s*), and thalamic reticular (*r*), as shown in **Figure 1A**. Depending on the populations, the connections between them can be excitatory, zero (not connected), or inhibitory ^27^. For each population *A*, the mean firing rate *Q*_*A*_, given a mean soma voltage *V*_*A*_, can be expressed with the nonlinear sigmoid function *Q*_*A*_ = *S*(*V*_*A*_) as in equation (1). The potential *V*_*A*_ is computed from the firing activity *ϕ*_*B*_ in other connected populations as follows:

**Figure 1.**
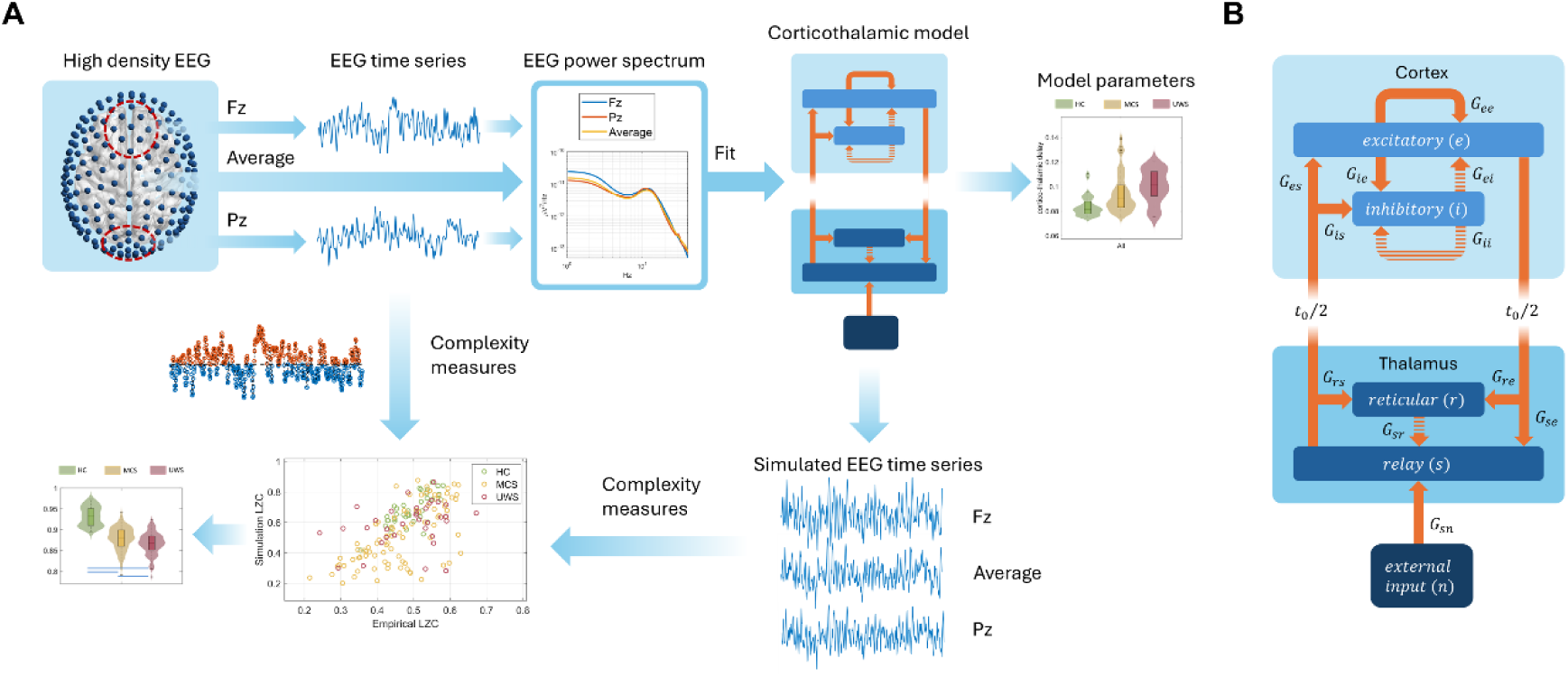
**A Protocol of the study.** High density EEG recordings from patients with prolonged DoC were used to estimate the parameters of a corticothalamic neural field model. To this end, the average power spectrum of 183 scalp channels, as well as the average from electrodes Fz, Pz and surrounding electrodes, were obtained. After fitting the model, time series were simulated for all subjects using the parameters obtained from the three power spectra. The Lempel-Ziv complexity of the empirical and simulated time series was obtained, their correlation was computed and their capacity to replicate group differences was assessed. **B Corticothalamic neural field model.** The cortex component consists of an excitatory population (e) and an inhibitory population (i); the thalamus component consists of a reticular population (r) and a relay population (s). The relay population receives external inputs denoted by n. The non-zero gains between each two populations are denoted by *G*_*ab*_. Inhibitory connections are shown in dashed lines. There is no time delay between populations within the same anatomical part, but a loop delay of *t*_0_ exists for any connection between cortex and thalamus.

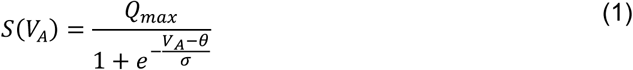

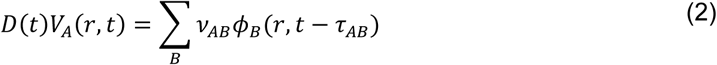

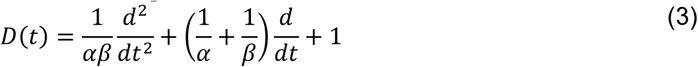

where *V*_*A*_(*r*, *t*) denotes the potential of population *A* at position *r* and time *t*, *θ* is the threshold that the soma voltage needs to exceed to produce an action potential, *Q*_*max*_ is the maximum of the sigmoid function, and *σ* is the variance of the threshold. The synaptic connection to population *A* from population *B* is *v*_*AB*_ = *N*_*AB*_ *s*_*AB*_, where *N*_*AB*_ is the mean number of synapses from *B* to *A*, and *s*_*AB*_ is the time-integrated response to a unit input signal. *ϕ*_*B*_(*r*, *t* − *τ*_*AB*_) is the mean firing arrival rate, allowing for a time delay *τ*_*AB*_ due to anatomical separation between populations. This means *τ*_*AB*_ is nonzero, equal to 1/2 of the corticothalamic propagation time (*t*_0_), only when *A* and *B* are cortical and thalamic populations respectively. α and β are the inverse rise and decay time of the cell-body potential produced by an impulse at a dendritic synapse. Thus equation (2) could be rewritten as a set of equations for each population separately, only taking into consideration the terms with nonzero parameters:

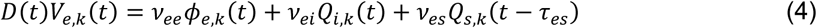

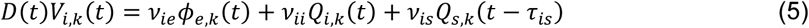

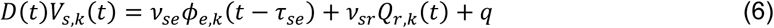

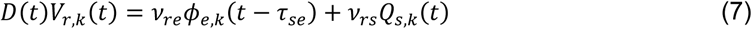

where *q* is a constant noise input to the thalamus relay population.

The firing rate produced by a population need to travel to other populations over long range white matter tracts. This spreading activity obeys the damped wave equation:

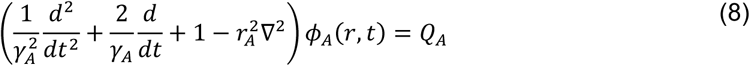

where *γ*_*A*_ refers to the cortical damping rate; 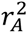 is the mean range.

These equations enable us to simulate the EEG-like time series. Since we are interested in the spectral features of the simulation, this could be done more efficiently by means of a transformation of the model into the frequency domain, as addressed in the original papers on the Robinson model ^27,59^. By setting all the derivatives to 0 in (3) and (8), we can obtain spatially uniform steady states and solve the steady state firing rate of the excitatory population 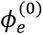:

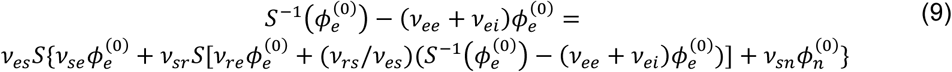

where *S*^−1^ denotes the inverse of the sigmoid function, 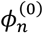 is the steady state component of the input stimulus. We can further approximate the brain’s activity as linear, *i.e*., first order, perturbation relative to a steady state. By linearizing equation (1) and keeping only the first order term in the Taylor expansion equation, we get:

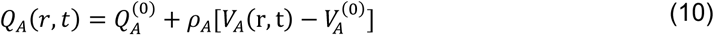

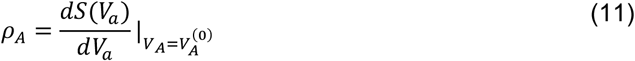

Next, applying a Fourier transform on the differential operator in equation (2), (3) and (8) yields the transfer function of *ϕ*_*e*_(*k*, *ω*) given a stimulus *ϕ*_*n*_(*k*, *ω*) of wave vector *k* and angular frequency *ω* ^29,59^:

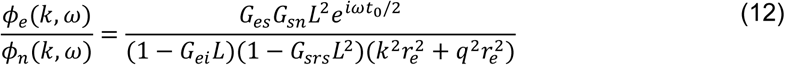

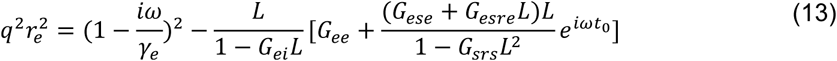

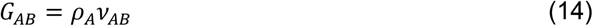

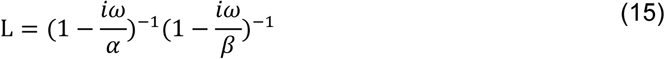

The gain *G*_*AB*_ denotes the differential response of population *A* per unit input from population *B*. And the gain loops are defined as: *G*_*ese*_ = *G*_*es*_ *G*_*se*_ for the excitatory corticothalamic loop through the relay population, *G*_*esre*_ = *G*_*es*_*G*_*sr*_ *G*_*re*_ for the inhibitory corticothalamic loop through both relay and reticular populations, *G*_*srs*_ = *G*_*sr*_ *G*_*rs*_ for the intrathalamic loop. The EEG power spectrum P(ω) can be calculated by integrating the squared magnitude of *ϕ*_*e*_(*k*, *ω*) achieved by integrating equation (9) over *k* ^27^:

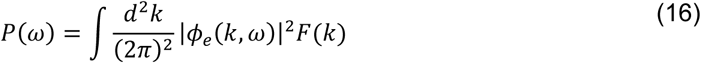

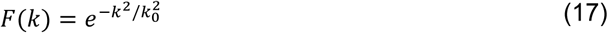

*F(k)* is a filter function that describes the volume conduction effect of the electrical activity of the cortex: the power in spatial modes with higher frequency (larger *k*) would be reduced due to the volume conduction of uncorrelated source activity. Here *k*_0_ = 10 *m*^−1^ is the state-independent low-pass cutoff ^60^.

Another important component in simulating the power spectrum is the electromyogram (EMG) component. The EMG signal is often observed at frequencies higher than 25 Hz in EEG recordings. It is caused by activity in pericranial muscles ^61,62^. Hence, an EMG component *P*_*EMG*_ is included in the model power spectrum ^63^:

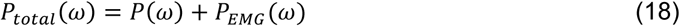

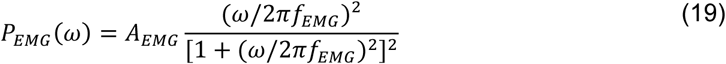

where *A*_*EMG*_ and *f*_*EMG*_ are the normalized EMG power and the characteristic EMG frequency, respectively. All the parameters and values can be found in Table 3 based on previous studies ^26,27,30^.

**Table 3.** Modelss parameter ranges and fixed values. The fixed parameters and ranges are based on previous literature ^26,27,30^.

| Symbol | Value or range | Initial Step | Unit |
| --- | --- | --- | --- |
| $\tau_{es}, \tau_{is}, \tau_{se}$ | 0.04 | - | s |
| $\gamma$ | 116 | - | s<sup>-1</sup> |
| $Q_{max}$ | 250 | - | s<sup>-1</sup> |
| $\theta$ | 15 | - | mV |
| $\sigma$ | 3.3 | - | mV |
| $r_e$ | 86 | - | mm |
| $k_0$ | 29 | - | m<sup>-1</sup> |
| $1/\alpha$ | 10-100 | 5 | ms |
| $1/\beta$ | 100-800 | 40 | ms |
| $G_{ee}$ | 0-20 | 0.4 | - |
| $G_{ei}$ | -40-0 | 0.4 | - |
| $G_{es}$ | 0-20 | 0.4 | - |
| $G_{se}$ | 0-20 | 0.4 | - |
| $G_{sr}$ | -40-0 | 0.4 | - |
| $G_{sn}$ | 0.8 | - | - |
| $G_{re}$ | 0-20 | 0.4 | - |
| $G_{rs}$ | 0-20 | 0.4 | - |
| $t_0$ | 0.075-0.14 | 0.005 | s |
| $G_{ese}$ | 0-40 | 1 | - |
| $G_{esre}$ | -40-0 | 1 | - |
| $G_{srs}$ | -5-0 | 0.2 | - |
| $P_{EMG}$ | 0-1 | 0.05 | mV |

The fitting process requires the “BrainTrak” library on MATLAB, which was developed as a method to analyze the sleep EEG signals and estimate corticothalamic model parameters for different brain activity in wake-sleep cycle ^28,64^. The code is available on GitHub: https://github.com/BrainDynamicsUSYD/braintrak. The model optimization is estimated by minimizing the chi-squared (*χ*^2^) error between the simulated and empirical power spectrum:

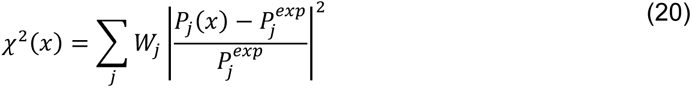

where the error is weighted by *W*_*j*_ = *f*_*j*_^−1^ for each frequency component *j* to compensate for the large number of sampling points at high frequencies compared to low frequencies, which results from the logarithmic transformation of the signal frequencies. Thus, the likelihood of the parameters is estimated by:

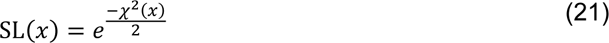

so that the likelihood is maximized to 1 when the error is 0 and is minimized to 0 as the error increases. The fitting starts with a set of parameters gained from previous fittings of healthy subjects’ resting state data ^28^. Then, a Markov Chain Monte Carlo sampling algorithm is used to characterize the posterior distribution of the parameters. At each step, a new random set of parameters is proposed by drawing a sample from a simple proposal distribution centered on the current parameter values. The new set of parameters is accepted if it yields a higher likelihood defined by equation (21). Otherwise, it is accepted with a probability equal to the ratio of the posterior probability of the new parameters to the posterior probability of the old parameters. This procedure will be repeated until there is no iterative change in the sampled probability distribution. The sequence of the accepted parameters in the random walk is referred to as the “chain”. After accepting 100 points on the chain, the parameter probability distribution is switched from an assumed one to the empirical one, which is then used to propose the subsequent step of the random walk. The first 5,000 points on the chain are discarded because they largely rely on the initial values. The total length of the preserved chain is 50,000 points. The final parameters are obtained by selecting the point on the chain that has the largest value of likelihood *L*(*x*). The probability distribution of each parameter indicates whether a good fit is achievable with a narrow or wide choice of parameters.

After fitting the power spectrum, we used the “NFTsim” software package, written in C++ and bundled with MATLAB routines ^65^ to generate time series given the fitted parameters. The main idea is to inversely solve equation (21) to get the firing response *Q*_*A*_ for each population *A* in the time domain. Afterwards, a neural field is created on a square domain with 144 grid cells in total, of which the spatially extended domain remains invariant throughout the simulation. The propagation process with time delay is constructed based on the physical length of the two dimensions in this field. The code for NFTsim is available on GitHub: https://github.com/BrainDynamicsUSYD/nftsim.

The power spectral density of the EEG signals used for model fitting was computed using the Welch method, with a segment length of 128 samples, an overlap of 100 samples, and a fast Fourier transform length of 4096. The PSD was computed for frequencies ranging from 1 to 40 Hz. We fitted 9 parameters for the spectrum fitting process: *G*_*ee*_, *G*_*ei*_, *G*_*ese*_, *G*_*esre*_, *G*_*srs*_, *⍺*, *β*, *t*_0_, *P*_*EMG*_, as indicated in equation (12) and (13), to be able to simulate the power spectrum. However, the process of simulating the time series requires the specific value of every gain between each pair of populations, which cannot be obtained directly from these 9 parameters, so we increased the fitted parameter number to 11: *G*_*ee*_, *G*_*ei*_, *G*_*es*_, *G*_*se*_, *G*_*sr*_, *G*_*rs*_, *G*_*re*_, α, β, *t*_0_, *P*_*EMG*_. The simulated time series had a length of 60 seconds, with a step width of 0.0005 seconds.

The fitting of the power spectrum was performed for the average of all 183 electrodes, as well as separately for the average values of Fz and Pz along with 8 immediately surrounding electrodes, respectively. Afterwards, the complexity of the empirical and simulated time series was estimated over the epochs by means of the LZC algorithm ^66^, the spectral entropy (SE) ^39^ and the permutation entropy (PE) in the theta band ^38^. The PE was computed in the theta band (PE_*θ*_) and with an embedding dimension of 3, due to previous literature indicating it is particularly effective at discriminating between UWS and MCS patients ^22,41^, see ^40^ for the specific implementation. Finally, the correlation between empirical and simulated measures was estimated by means of Spearman’s coefficient. **Figure *1***B shows a schematic description of the main steps of the study.

### Statistical analyses

The values of *G*_*ee*_, *G*_*ei*_, *G*_*ese*_, *G*_*esre*_, *G*_*srs*_, *⍺*, *β*, *t*_0_, *P*_*EMG*_ and empirical and simulated LZC, PE and SE were compared among groups and electrodes by means of non-parametric statistical tests. Kruskall-Wallis tests were used to detect global interactions between the three groups (HC, MCS and UWS), and if interactions were found, Mann-Whitney *U*-tests were performed to determine between-group differences. To account for multiple comparisons, a false discovery rate (FDR) correction was applied ^67^. The significance level was set at α=0.05. The correlation strength was interpreted following conventional thresholds: high (|*r*| > 0.6), moderate (0.6 ≥ |*r*| > 0.4) and low (|*r*| ≤ 0.4). Statistical analyses and signal processing were performed using MATLAB^®^ (version R2024b Mathworks, Natick, MA).

## Results

### Model parameters

Examples of power spectrum fitting for a representative subject from each group are shown in **Figure *2***A. The model successfully fit the spectra across all groups with low chi-square error, and no significant differences in fitting error were observed between groups, as shown in **Figure *2***B. The Kruskal-Wallis tests revealed global interactions between the three groups for both the Liège and Paris datasets in model parameters, specifically between the control and patient groups. No significant differences were found between the frontal and parietal channels, so the subregional analyses were not presented in the main results but can be found in supplementary Tables S1-S2, and Figures S1-S2. No significant correlation was found between the model’s parameters and the TSO or CRS-R index (Table S3, S4). The parameters that exhibited significant group effects in both datasets were the following (see Figure 3; the statistics and *p*-values can be found in Table 4 and Table 5 for the Liège and Paris datasets, respectively) *G*_*ese*_, excitatory corticothalamic loop gain, which was lower in patients; 2) *G*_*esre*_, the inhibitory corticothalamic loop gain, which was higher in patients; 3) *G*_*srs*_, the intrathalamic loop gain, which was higher in patients; 4) *t*_0_, the delay between cortical and thalamic populations, which was higher in patients; and 5) *P*_*EMG*_, the EMG component, which was higher in patients.

**Figure 2.**
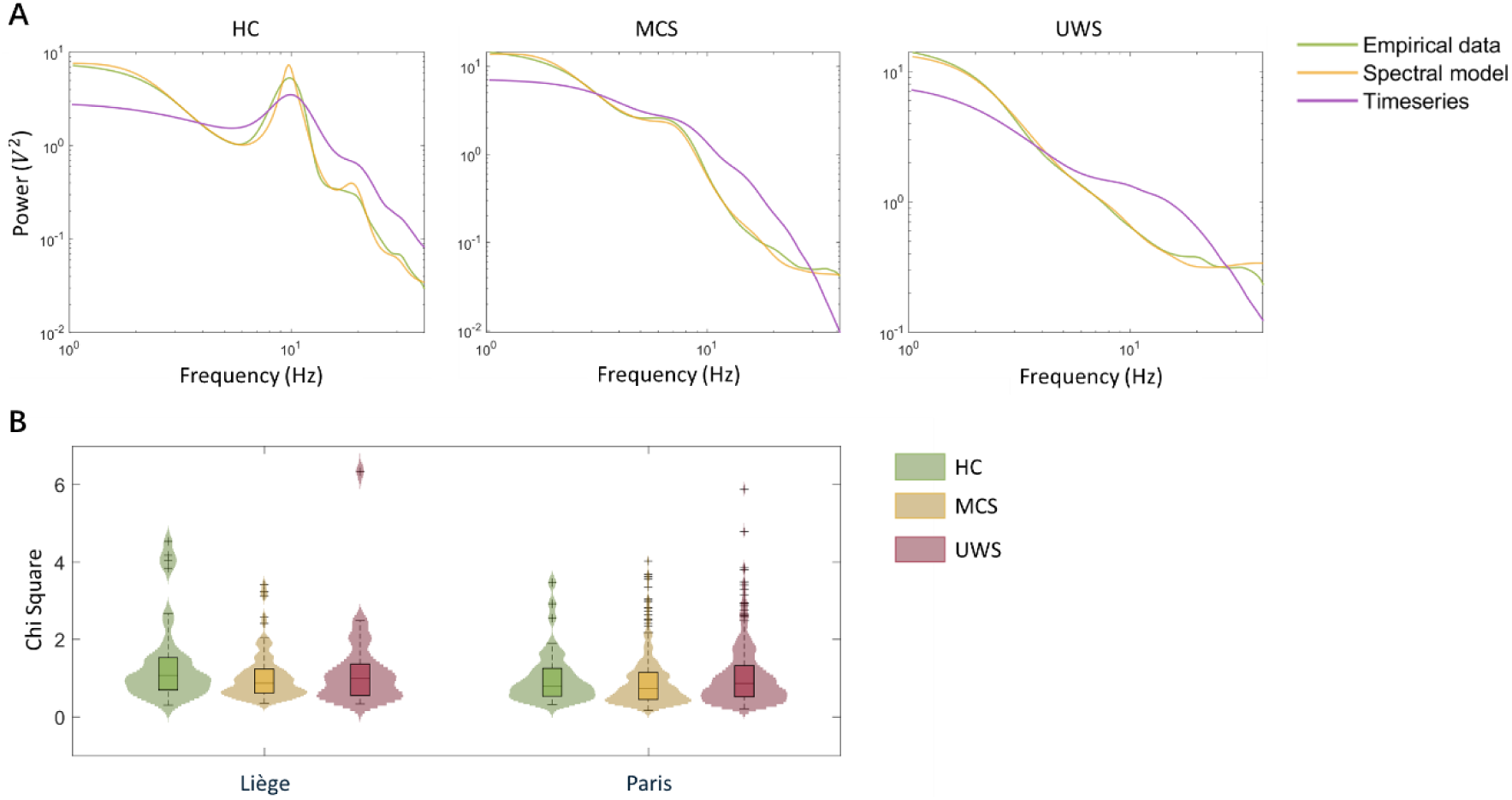
**A Power spectra for one representative subject from each group.** The power spectra were plotted on a log-log scale over 1-40 Hz. The green line represents the empirical data used as input to the model, the yellow line shows the model’s output, and the purple line displays the power spectrum of a simulated time series based on the model’s parameter settings. This illustrates how well the model captures the key spectral features of the recorded data across groups. **B Chi-squared error between the empirical and simulated power spectra**. The fitting errors are consistently low across all subject groups and datasets (Liège and Paris), and no statistically significant difference in error was found between groups.

**Figure 3.**
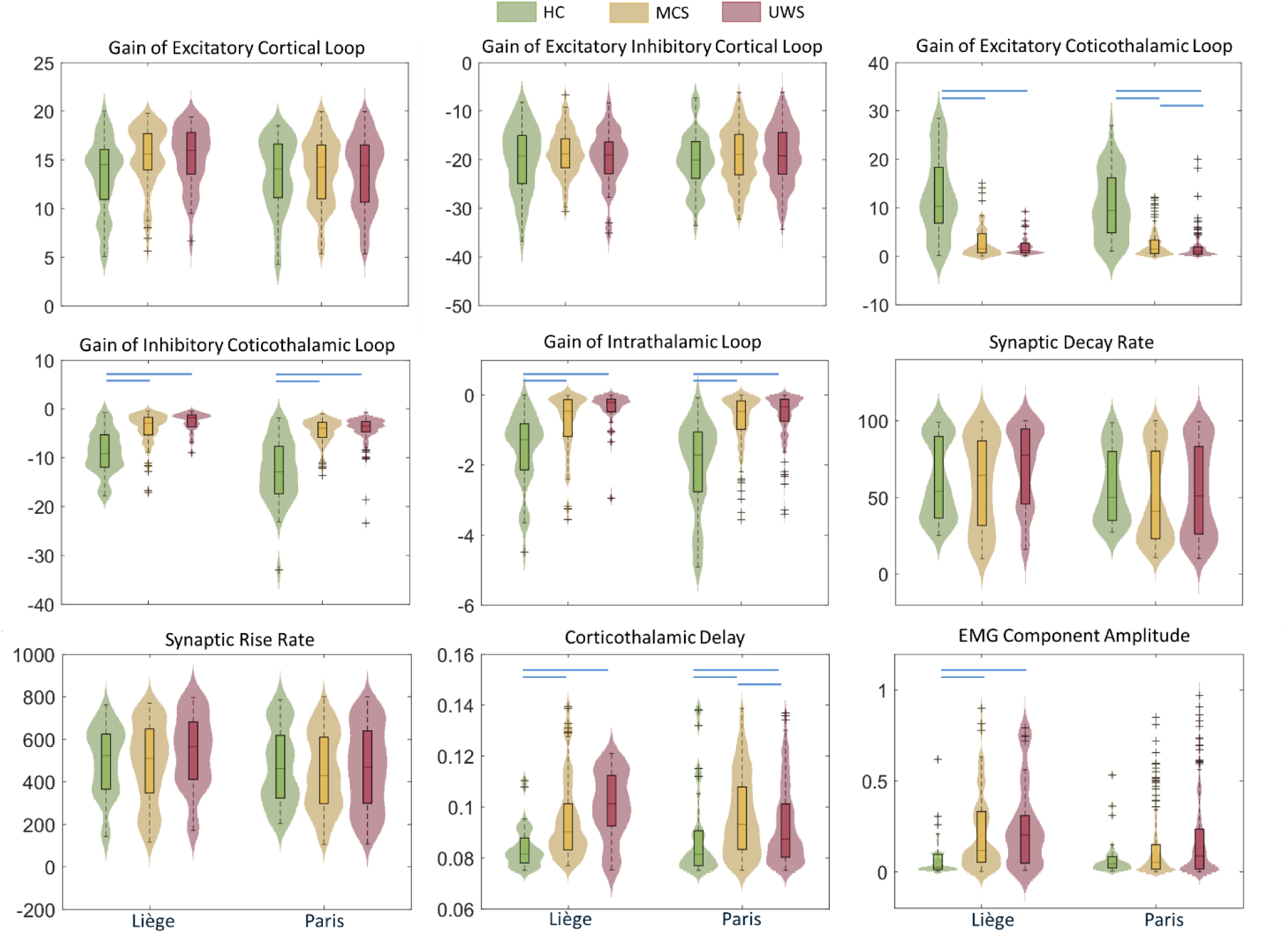
Comparisons of the model parameters for the three subject groups in two datasets. Significant differences were found between HC and DoC patients in both datasets in the following parameters: 1) *G*_*ese*_, the gain of the excitatory corticothalamic loop was lower in DoC patients; 2) *G*_*esre*_, the gain of the inhibitory corticothalamic loop is higher in DoC patients; 3) *G*_*srs*_, the gain of the intrathalamic loop was higher in DoC patients; 4) *t*_0_, the delay between cortical and thalamic regions was longer in DoC patients; 5) the EMG component amplitude was larger in DoC patients. Moreover, three parameters differed between MCS and UWS patient groups in the Paris dataset only: 1) *G*_*esre*_ was larger in the MCS group than in UWS group; 2) *t*_0_ was larger in MCS group than in the UWS group. Blue lines indicate a significance level of p<0.05.

**Table 4.**
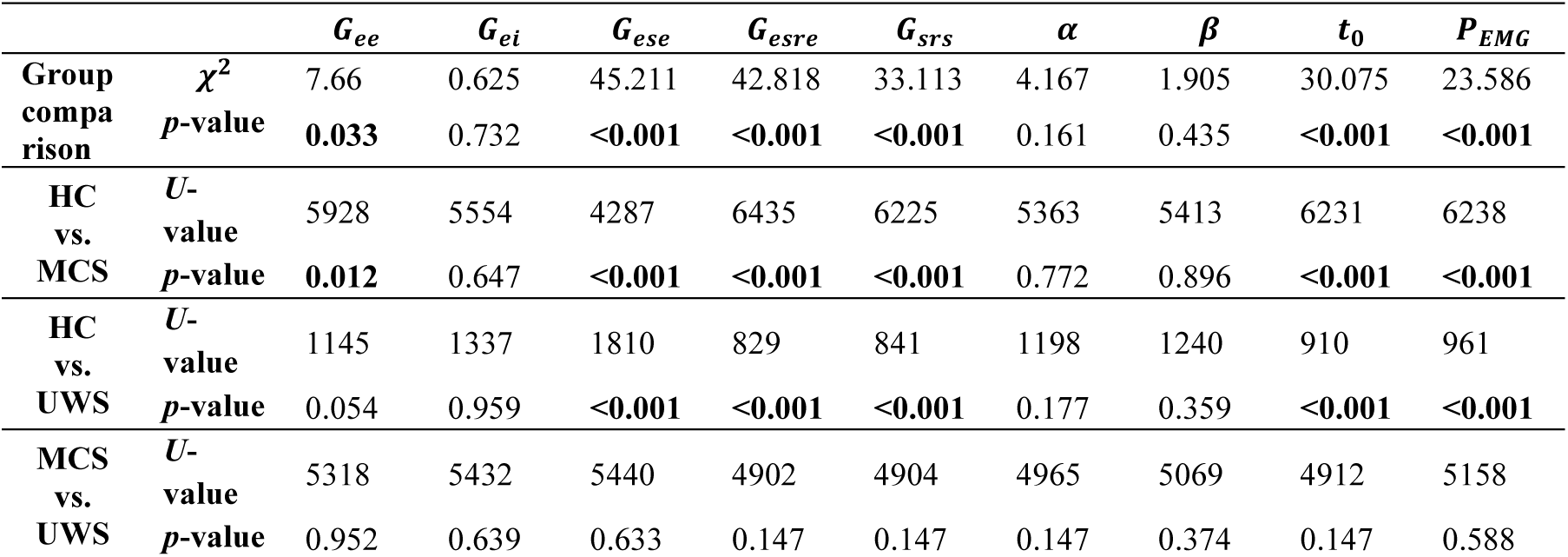
Statistical comparisons of the group (Kruskal-Wallis test) and pairwise (Mann-Whitney *U*-test) interactions between groups (UWS, MCS, HC) of the model parameters in the Liège dataset. FDR corrected for the number of comparisons. UWS: unresponsive wakefulness syndrome. MCS: minimally conscious state. HC: healthy controls.

**Table 5.** Statistical comparisons of the group (Kruskal-Wallis test) and pairwise (Mann-Whitney *U*-test) interactions between groups (UWS, MCS, HC) of the model parameters in the Paris dataset. FDR corrected for the number of comparisons. UWS: unresponsive wakefulness syndrome. MCS: minimally conscious state. HC: healthy controls.

| | | $G_{ee}$ | $G_{ei}$ | $G_{ese}$ | $G_{esre}$ | $G_{srs}$ | $\alpha$ | $\beta$ | $t_0$ | $P_{EMG}$ |
| --- | --- | --- | --- | --- | --- | --- | --- | --- | --- | --- |
| **Group comparison** | $\chi^2$ | 4.712 | 2.505 | 68.567 | 63.89 | 50.739 | 4.946 | 3.391 | 22.072 | 11.592 |
|  | *p*-value | 0.720 | 0.257 | **<0.001** | **<0.001** | **<0.001** | 0.213 | 0.678 | **<0.001** | 0.065 |
| **HC vs. MCS** | *U*-value | 37143 | 37296 | 33579 | 40720 | 40341 | 36110 | 36866 | 39049 | 36903 |
|  | *p*-value | 0.611 | 0.568 | **<0.001** | **<0.001** | **<0.001** | 0.119 | 0.611 | **<0.001** | 0.611 |
| **HC vs. UWS** | *U*-value | 3873 | 3323 | 6437 | 1342 | 1420 | 4083 | 3904 | 3034 | 3201 |
|  | *p*-value | 0.941 | 0.168 | **<0.001** | **<0.001** | **<0.001** | 0.614 | 0.941 | **0.031** | 0.090 |
| **MCS vs. UWS** | *U*-value | 4684 | 5087 | 5409 | 4968 | 4818 | 4850 | 5154 | 5474 | 4578 |
|  | *p*-value | 0.455 | 0.267 | **0.043** | 0.061 | 0.070 | 0.249 | 0.420 | **0.043** | 0.061 |

Subsequent pairwise comparisons confirmed that these differences were statistically significant between the HC and both patient groups individually for both datasets (MCS and UWS, *p* <0.001, Mann-Whitney *U* test; see Table 4 and Table 5 for the test statistics and *p*-values for the Liège and Paris datasets, respectively). Moreover, in the Liège dataset, *G*_*ee*_, the gain between cortical excitatory populations, was lower in the control group than in the MCS group (*p* = 0.012); and *P*_*EMG*_, the EMG component, was higher in patients than in the MCS and UWS group, with both p-values < 0.001.

Mann-Whitney *U*-tests revealed significant differences between the MCS and UWS groups in the Paris dataset. The gain of the excitatory corticothalamic loop (*G*_*ese*_) was higher in the MCS group than in the UWS group (*p* = 0.043). Similarly, the corticothalamic delay (*t*_0_) was longer in the MCS group, with a *p*-value of 0.043. No difference was found between MCS and UWS groups in Liège dataset. Nevertheless, the median value for *G*_*ese*_ was higher in the MCS group, consistent with the Paris result. The trend of *t*_0_ was not consistent across datasets.

### Correlation of complexity features between empirical and simulated data

We computed LZC, SE, and PE_*θ*_ for both the empirical and simulated data and evaluated the correlation between the two parameter sets within each group. We first examined the difference of complexity measures between frontal and parietal channels across subject groups but found no significant effect (Table S5). Therefore, we report only the results using the 183-channel recordings, with the full statistics results in Table S6.

As shown in **Figure *4***, in the Liège empirical dataset, no significant group difference in LZC was found, but the Paris empirical dataset showed a significant group effect (*p* < 0.001), with post hoc tests revealing that LZC of HC was significantly higher than that of MCS (*p* < 0.001) and UWS (*p* < 0.001). Similarly, in Liège simulated time series, we found that the LZC of the HC group was higher than the MCS group (*p* < 0.001), and that the LZC of UWS was higher than MCS (*p* = 0.008). In the simulation for the Paris dataset, we found that the LZC of the HC group was higher than the MCS (*p* < 0.001) and UWS (*p* < 0.001). Additionally, we found significant correlation between the LZC of empirical and simulated data, as visualized in **Figure *5***. In the Paris dataset, the correlation coefficients were 0.655 (*p* < 0.001) for HC, 0.449 (*p* < 0.001) for MCS, and 0.254 (*p* < 0.001) for UWS. In the Liège dataset, the correlations were 0.717 (*p* < 0.001) for HC, 0.542 (*p* < 0.001) for MCS, and 0.378 (*p* = 0.027) for UWS. For all combined data in two datasets the correlation coefficient was 0.458 (p<0.001). Statistical results are shown in Table 6.

**Figure 4.**
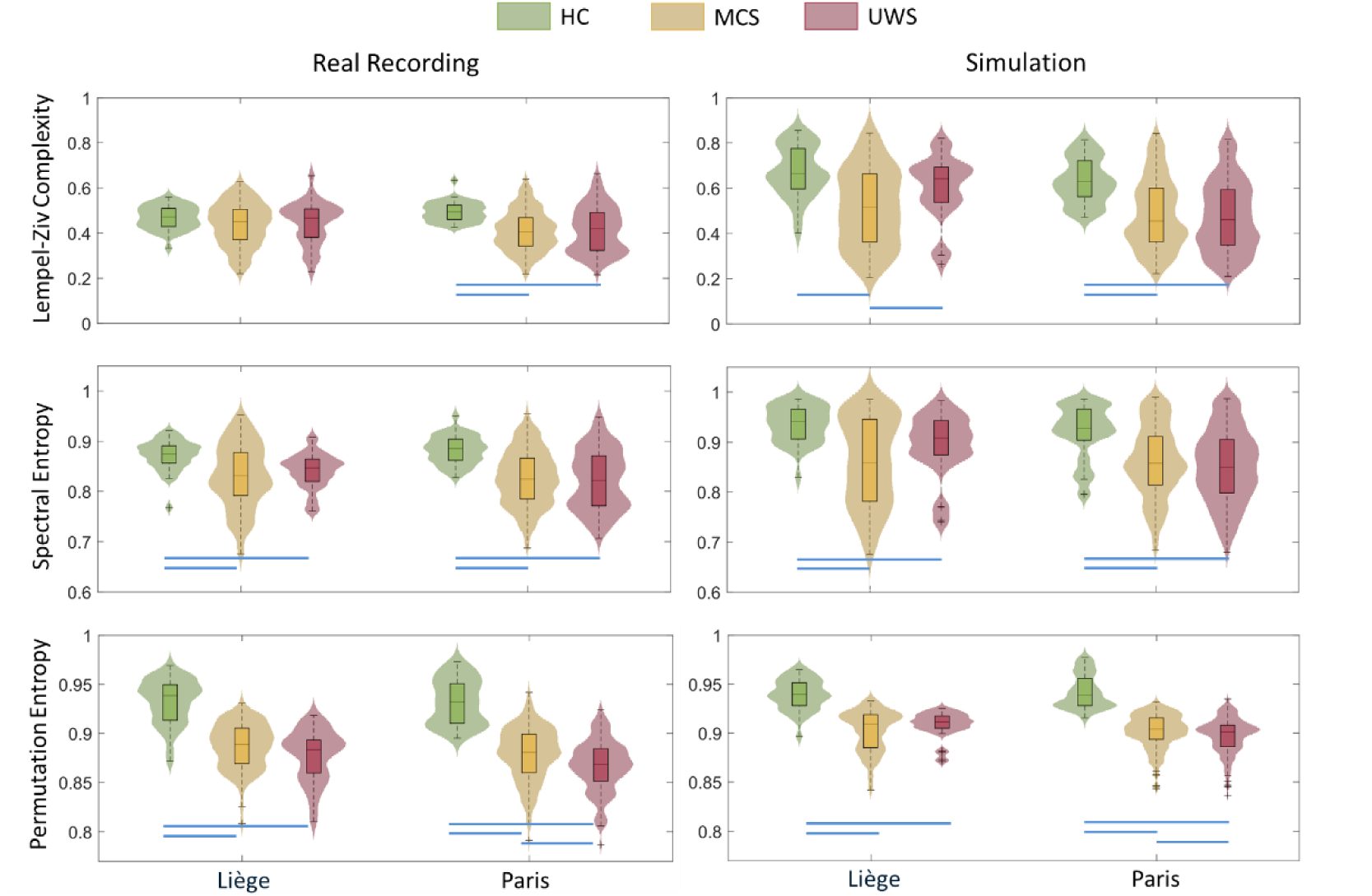
Comparison of complexity measures of real and simulated time series across subject groups in two datasets. In the Liège dataset, the SE and PE of the time series were significantly higher in the HC group than in the two patient groups in both real and simulated data. In the Paris dataset, the LZC, SE and PE of the time series were significantly higher in the HC group than in the two patient groups in both real and simulated data; in addition, the PE was found to be significantly higher in the MCS group than in the UWS group. The simulated time series reproduced this overall pattern, with one exception: in the Liège dataset, the LZC of the MCS group was lower than that of both the HC and UWS groups. Blue lines indicate a significance level of p<0.05.

**Figure 5.**
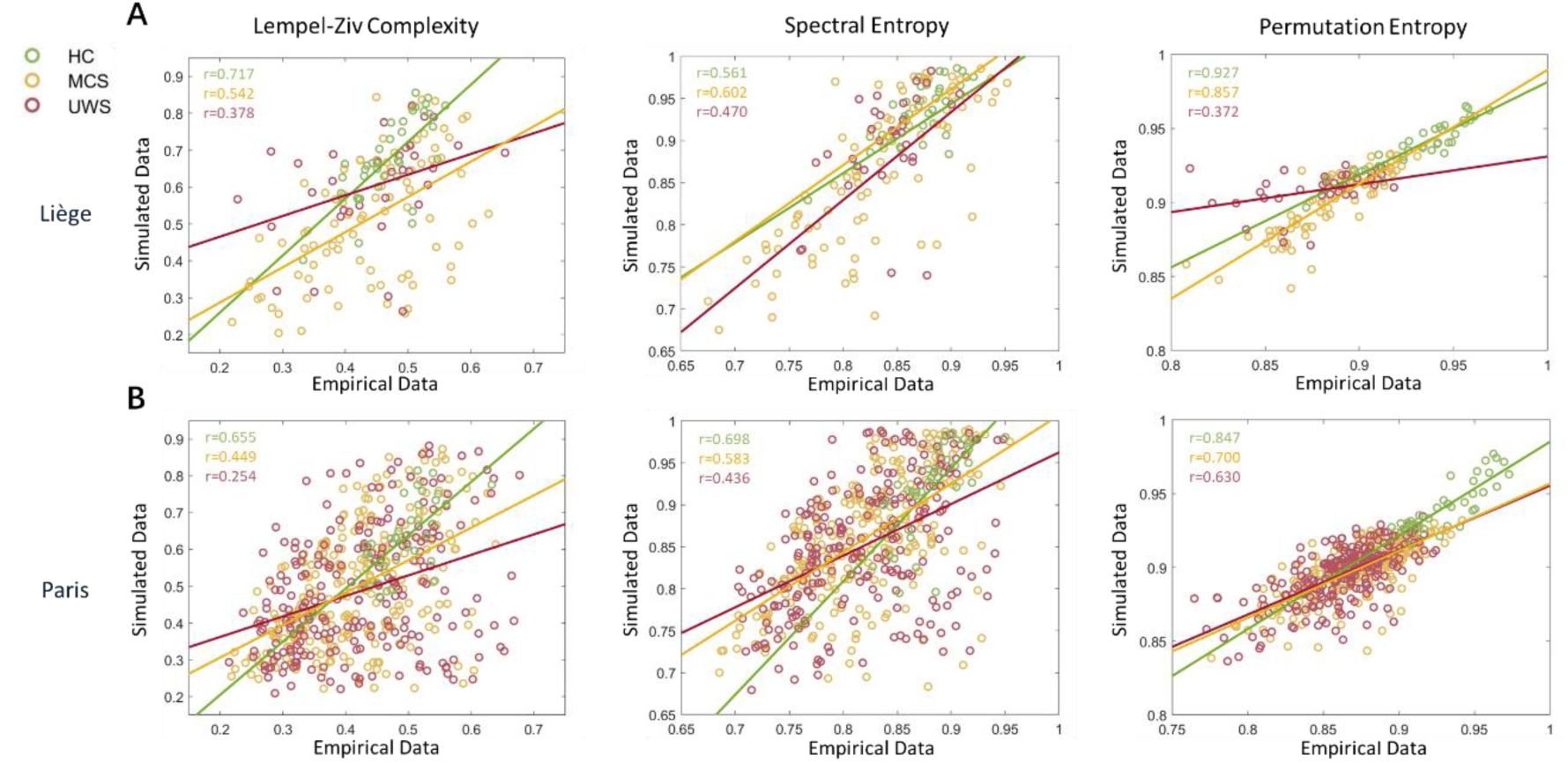
Correlation of the complexity measures between empirical and simulated data in the two datasets. Panel A and B show separately the correlation plots for the Liège and Paris datasets, including Lempel-Ziv complexity (LZC), spectral entropy (SE), and theta-band permutation entropy (*PE*_*θ*_). In general, *PE*_*θ*_ showed the strongest correspondence between real and simulated data, and correlations were consistently higher in the healthy control group compared to patient groups.

**Table 6.**
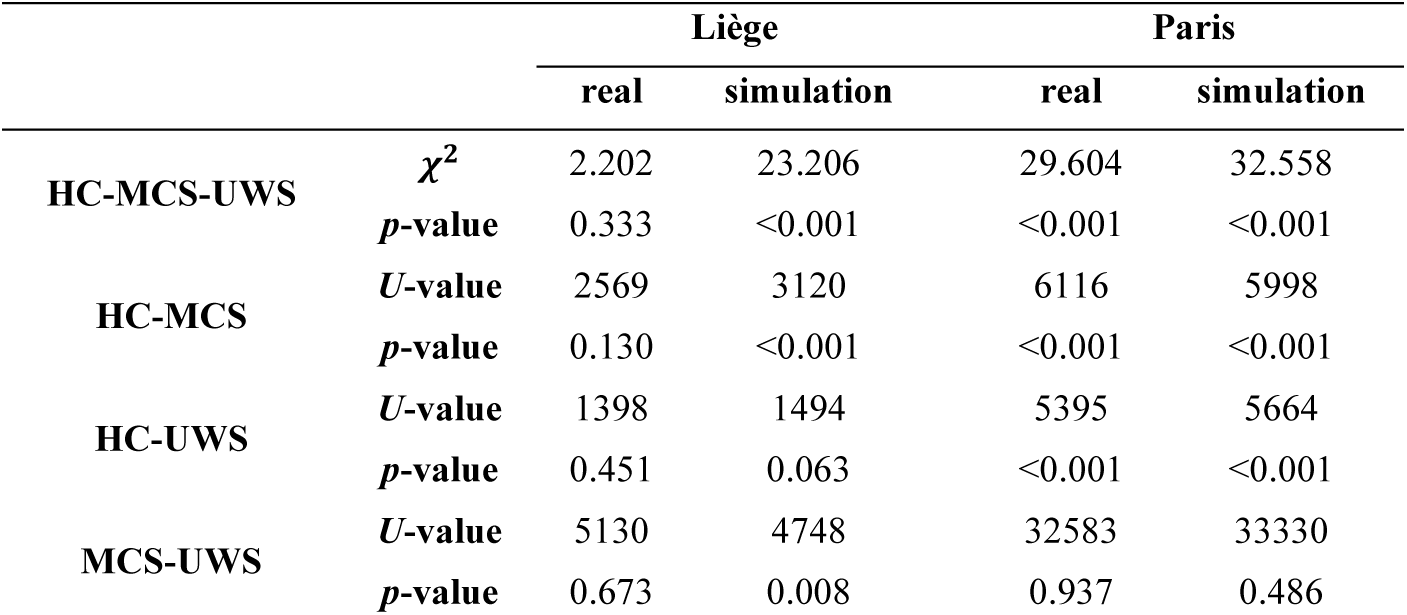
Statistical comparisons of the group and pairwise interactions between groups (UWS, MCS, HC) of the Lempel-Ziv Complexity (LZC) of the real and simulated time series in two datasets. Global interactions were evaluated with Kruskal-Wallis tests, while pairwise interactions were evaluated with Mann-Whitney *U*-tests.

Spectral entropy (SE) was also computed for both empirical and simulated data and the results are shown in **Figure *4***. In both the Liège and Paris datasets, SE differed significantly across subject groups (Liège: *p* < 0.001; Paris: *p* < 0.001). Post-hoc comparisons showed that SE was significantly higher in the HC group than in both patient groups in both datasets (both datasets *p* < 0.001). The same pattern was observed in the simulated data: in Liège, SE was higher in HC compared to MCS (*p* < 0.001) and UWS (*p* = 0.003); in Paris, HC also showed higher SE than MCS (*p* < 0.001) and UWS (*p* < 0.001). However, no significant difference was found between the MCS and UWS groups in either dataset. We also found a moderate to high correlation between the SE of real and simulated data in both datasets (**Figure *5***): ρ = 0.698 for HC, ρ = 0.583 for MCS and ρ = 0.436 for UWS in the Paris dataset (all *p* < 0.001); in the Liège dataset the correlations were ρ = 0.561 (p<0.001) for HC, ρ = 0.602 (p<0.001) for MCS and ρ = 0.470 (p=0.005) for UWS. For all combined data in two datasets the correlation coefficient was 0.613 (p<0.001). Statistical results can be found in Table 7.

**Table 7.** Statistical comparisons of the group and pairwise interactions between groups (UWS, MCS, HC) of the Spectral Entropy (SE) of the real and simulated time series in two datasets. Global interactions were evaluated with Kruskal-Wallis tests, while pairwise interactions were evaluated with Mann-Whitney *U*-tests.

|  |  | Liège |  | Paris |  |
| --- | --- | --- | --- | --- | --- |
|  |  | real | simulation | real | simulation |
| HC-MCS-UWS | $\chi^2$ | 17.015 | 20.633 | 37.079 | 27.283 |
|  | *p*-value | <0.001 | <0.001 | <0.001 | <0.001 |
| HC-MCS | *U*-value | 2979 | 3075 | 6182 | 5723 |
|  | *p*-value | <0.001 | <0.001 | <0.001 | <0.001 |
| HC-UWS | *U*-value | 2999 | 1599 | 5727 | 5562 |
|  | *p*-value | <0.001 | 0.003 | <0.001 | <0.001 |
| MCS-UWS | *U*-value | 7264.5 | 4928 | 29375 | 29048 |
|  | *p*-value | 0.296 | 0.109 | 0.406 | 0.242 |

Finally, we examined the theta band permutation entropy (PE_*θ*_) of the time series (**Figure *4***). In both datasets, Mann-Whitney *U* tests revealed that the PE_*θ*_ was significantly higher in healthy controls than in the patients (Paris: *p* < 0.001; Liège: *p* < 0.001; all pairwise control vs. patient group tests with *p* < 0.001). Notably, in the Paris dataset, the PE_*θ*_ of the MCS group was higher than that of the UWS group (*p* < 0.001). Simulated data mirrored these results, with significant group effects in both datasets (Paris: *p* < 0.001; Liège: *p* < 0.001), and post-hoc comparisons confirmed higher PE_*θ*_ values in HC than in the patient groups (all *p* < 0.001). In the Paris simulated data, the MCS group had higher PE_*θ*_ than the UWS group (*p* = 0.018). A high correlation was found between the PE_*θ*_ of real and simulated data in Paris dataset (**Figure *5***): 0.847 for HC, 0.700 for MCS and 0.630 for UWS, (all *p* < 0.001); In the Liège dataset, the correlation of PE_*θ*_ of real and simulated data was high for HC (ρ = 0.927, *p* < 0.001) and MCS (ρ = 0.857, *p* < 0.001), and lower for UWS (ρ = 0.372, *p* = 0.030). For all combined data in the two datasets the correlation coefficient was 0.803 (p<0.001). Detailed statistics results are provided in Table 8.

**Table 8.**
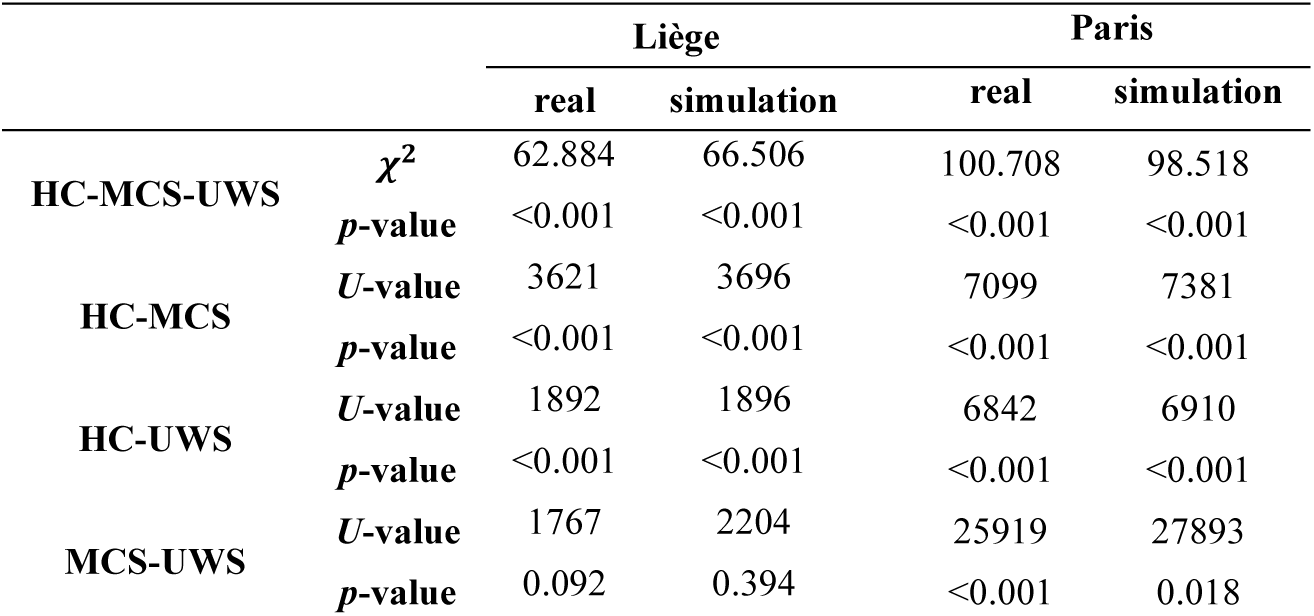
Statistical comparisons of the group and pairwise interactions between groups (UWS, MCS, HC) of the Permutation Entropy (PE) of the real and simulated time series in two datasets. Global interactions were evaluated with Kruskal-Wallis tests, while pairwise interactions were evaluated with Mann-Whitney *U*-tests.

|  |  | Liège |  | Paris |  |
| --- | --- | --- | --- | --- | --- |
|  |  | real | simulation | real | simulation |
| HC-MCS-UWS | $\chi^2$ | 62.884 | 66.506 | 100.708 | 98.518 |
| | $p$-value | <0.001 | <0.001 | <0.001 | <0.001 |
| HC-MCS | $U$-value | 3621 | 3696 | 7099 | 7381 |
| | $p$-value | <0.001 | <0.001 | <0.001 | <0.001 |
| HC-UWS | $U$-value | 1892 | 1896 | 6842 | 6910 |
| | $p$-value | <0.001 | <0.001 | <0.001 | <0.001 |
| MCS-UWS | $U$-value | 1767 | 2204 | 25919 | 27893 |
| | $p$-value | 0.092 | 0.394 | <0.001 | 0.018 |

See supplementary material Figure S3 and Figure S4 for the comparisons between frontal and parietal channels, as well as the correlation analyses for the full data in Table S7. We found that the complexity was higher in the TSO>1 group than in the TSO<1 group, which is significantly shown in all three measures in the Paris dataset and the PE in the Liège dataset (Figure S5 and Table S8). Low correlation was found between the CRS-R index and SE and PE values in the Paris dataset (Table S9).

## Discussion

Given the presumed importance of corticothalamic dynamics in consciousness, in this study we employed a corticothalamic neural field model to describe and simulate EEG recordings across HC and patients with DoC. To this end, we utilized two large, independent high-density EEG datasets, combining resting-state and LG recordings, which included a total of 277 UWS, 340 MCS, and 74 controls. We aimed to identify specific alterations in cortico-thalamic circuits that distinguish between the brain states of healthy individuals and patients, as well as between awareness and unawareness.

### Corticothalamic parameters and the role of thalamocortical loops

The results demonstrated that the corticothalamic model accurately captured key spectral features across groups. It was able to reproduce the position and amplitude of the alpha peak in the HC subjects, the prominent theta band activity in the MCS patients, and the delta-dominated power spectrum in UWS patients. Previous studies showed its ability to simulate healthy resting state activity and sleep spindles ^30^, periodic spikes during epileptic seizures ^31^. It has been observed that, in the case of complete thalamocortical deafferentation, spectral peaks can be absent altogether ^68^, which was the case in some UWS patients, and the model was able to reliably replicate the absence of periodic components in the spectrum. Our findings further validate the model’s applicability in a variety of scenarios by showing its ability to simulate brain states of healthy individuals and brain-injured patients in two independent datasets (Liège and Paris), where the latter was able to distinguish between UWS and MCS, extending previous findings ^33^. We thus confirmed the model’s ability to fit and simulate the comparatively smoother PSD curve of DoC patients. The model’s high goodness of fit across all groups further supports its suitability for estimating the connectivity between the neuronal populations it incorporates, as reflected in its parameter values.

The variation in model parameters across groups partially aligns with the findings of Assadzadeh et al. ^33^. Although we did not replicate the group effect on synaptic rise or decay rates, we observed similar variance in corticothalamic loop parameters. The physiological interpretation of these parameter changes is consistent with the mesocircuit hypothesis mechanisms in the context of DoC. Specifically, in the patient group, we noted a lower absolute value for the gain of the corticothalamic excitatory and inhibitory loop, along with a longer delay between cortical and thalamic populations. This finding supports the mesocircuit hypothesis, which posits that disrupted EEG patterns in DoC patients are due to impairments in corticothalamic connectivity ^8^. This hypothesis has been substantiated by various studies employing different methodologies, including MRI ^69^, PET ^70^, EEG ^21^, and spectral dynamic causal modelling on MRI data ^71^. The results are also in agreement with those observed using the same model on the EEG of patients experiencing acute coma within the first 48 hours after cardiac arrest, in which patients with poor outcome showed increased corticothalamic inhibition, and higher delay compared to those with good outcome ^72^ Our study adds to the literature by further supporting this theory from the perspective of biophysically informed EEG modelling.

Beyond the differences observed between patients and healthy controls, the model also revealed a subtler, yet meaningful distinction between MCS and UWS patients in the Paris dataset, which was not previously reported by Assadzadeh et al. ^33^. Specifically, in the Paris dataset, we found that the gain of the excitatory corticothalamic loop was lower in UWS compared to MCS. This finding aligns with previous studies. For example, Magrassi et al. ^73^ reported increased activity of medial thalamic neurons and stronger phase coupling between thalamic discharges and cortical EEG signals in MCS patients compared to those with UWS, based on microelectrode recordings. Similarly, MRI studies have shown that thalamocortical pathways connecting the thalamus with the dorsal and posterior regions of the posteromedial cortex exhibit less white matter integrity in UWS patients than in MCS patients ^74,75^. We thus propose that the higher corticothalamic gain in the model may reflect a more preserved thalamocortical interaction in MCS. Considering the proven accuracy of the model in replicating spectral features after fitting, this finding would be in line with a previous study, which showed that a higher corticothalamic integrity, corresponding to certain EEG spectral features such as theta and beta peaks, is also indicative of an increased likelihood of recovery of consciousness in acute DoC patients ^9^.

Interestingly, we found that corticothalamic delay was shorter in UWS than in MCS. One possible interpretation is that shorter delays in UWS may reflect a breakdown of refined thalamocortical processing, leading to fast, asynaptic, and hyperexcitable physiological activity that is incapable of supporting consciousness. This aligns with evidence showing UWS patients have lower perturbational complexity index values (PCI) associated with stereotypical responses to TMS-EEG ^37^, altered fMRI signal propagation ^76^, and increase in thalamic hemodynamic response found in UWS patients ^77^. In contrast, the longer delay in MCS could indicate a partially preserved though potentially slowed thalamocortical circuit, which would allow for some conscious processing in the MCS patients, but the structural damage, such as axonal injury or demyelination ^17,78^, could lead to slowed signal transmission along thalamocortical fibers, thereby increasing effective delays.

In addition to the main findings, we observed that the EMG component was consistently larger in patients across the Liège dataset; Since the EMG component is introduced to account for non-neural, high-frequency noise, this pattern could reflect increased movement artifacts associated with patients’ physiological states. However, this interpretation should be approached with caution. The EMG term primarily serves as a mathematical ‘patch’, compensating for spectral features that are not well captured by the neural field model. Therefore, its variation may simply reflect a model mismatch in fitting the high-frequency range, rather than a meaningful physiological distinction between groups.

Contrary to our initial hypothesis, we found no significant differences between frontal and parietal regions across all the measures examined, including model parameters and signal complexity. This might be due to the fact that the corticothalamic model addressed different aspects of the EEG data. For instance, several investigations have suggested that parietal regions may be more sensitive than frontal areas in distinguishing MCS from UWS, particularly through time-varying gamma phase synchronization, mutual information, participation coefficient, and clustering coefficient ^40,54,79^. However, these metrics primarily emphasize functional connectivity and network-level integration, whereas our modelling approach focuses on reproducing the spectral power of EEG signals. As such, the lack of observed regional differences in our results may reflect a mismatch between the targeted neural features: power spectrum fitting may not be sufficiently sensitive to capture the dynamic inter-regional interactions that characterize conscious state differentiation.

### EEG complexity analyses

Having established that the model’s parameter space can distinguish clinical groups, we further analyzed the signal complexity of the real and simulated data, providing the link between the abstract parameters and measurable neurophysiological output. When simulating synthetic EEG signals with the fitted parameters, the model was able to replicate, to an extent, the features of the empirical recordings. We observed that certain spectral features, such as alpha and beta band oscillations, are relatively well replicated. These findings suggest that the time series simulations can effectively mimic specific characteristics of real EEG data. In addition, we found that the permutation entropy in the theta band more effectively discriminates between groups compared to Lempel-Ziv complexity (LZC) and spectral entropy (SE), both in real and simulated data. In the Paris dataset, the theta-band permutation entropy (PE_*θ*_) was significantly lower in the UWS patient group than the MCS patient group, consistently in the real and simulated EEG. This result is in agreement with Sitt et al. ^20^, where it was demonstrated that lower-frequency band complexity is a more robust and sensitive marker for distinguishing levels of consciousness in DoC patients, exhibiting a larger mean difference between groups and smaller within-group variability.

Additionally, the complexity measures of the simulated data showed a moderate to high correlation with those of real data: in all three measures, the correlations were higher for the healthy controls than for the patients. This discrepancy may reflect the larger inter-individual variability and spectral abnormalities found in pathological brain states, which challenged the model’s capacity to accurately capture their underlying dynamics. Among the metrics, the permutation entropy exhibited the strongest correspondence between modalities. This might also be due to the symbolic transformation used in this method, designed to enhance the SNR of the metric and the relatively low within-group variance of this measure, further supporting its reliability as a measure of EEG signal complexity in consciousness studies. The previously reported high diagnostic accuracy in discriminating UWS and EMCS patients provided by PE_*θ*_ in empirical recordings ^20,22^ is a key motivating factor in the potential use of the model not only as a biomarker for diagnostic purposes, but also for gaining mechanistic insight into each patient’s specific neural impairments. Furthermore, the model allows estimation of how close a patient’s brain dynamics are to criticality, a regime thought to support flexible, integrated information processing and consciousness ^80^. Deviations from criticality are a commonly observed feature of the brain dynamics in DoC^81,82^, making it a valuable target for intervention. By identifying whether a patient’s brain is in a subcritical or supercritical state, the model can help guide personalized therapeutic strategies, such as enhancing excitability in subcritical dynamics or stabilizing activity in supercritical ones.

In both the model parameter analysis and the time series complexity results, we observed that the difference between MCS and UWS patients was more pronounced in the Paris dataset than in the Liège dataset. It is essential to note that these datasets differ in their experimental design: the Liège dataset consists of resting-state recordings, whereas the Paris dataset utilizes data from a local–global auditory paradigm, which is treated here as a pseudo-resting state. Previous studies have demonstrated that the local-global paradigm can serve as a marker of conscious processing, with the global effect more frequently observed in MCS patients than in those with UWS ^40,53^. One possible explanation for our results is that task-free, resting state recordings, may fail to detect subtle residual signs of consciousness. In contrast, the local-global paradigm actively engages hierarchical processing, which might allow for the detection of brain responses more directly linked to conscious awareness and cognition, thus providing greater sensitivity in distinguishing between states of consciousness.

### Limitations and future directions

One limitation of this study is that we used scalp EEG recordings for the power spectrum fitting. The uncertainty of the original source of the activity recorded around the Fz and Pz electrodes might have hindered the detection of effects involving frontal and parietal areas that are central to dominant theories of consciousness, such as the GNW and the posterior “hot zone” IIT hypotheses ^43,83^. These spatial constraints are especially relevant for paradigms like the local-global task, where fronto-parietal dynamics play a critical role in conscious processing. In addition, we used a common average reference scheme, which, while commonly applied, can obscure localized neural activity by averaging signals across the entire scalp. More spatially specific referencing methods, such as current source density (CSD) could potentially provide better differentiation between nearby cortical sources ^84^. While source-reconstructed EEG could offer improved spatial mapping and better localization of underlying cortical sources, this was not feasible in the present study. Most patients lacked high-resolution T1-weighted MRI scans, and the substantial heterogeneity of brain lesions across individuals made it inappropriate to rely on standard anatomical templates, which could introduce significant spatial inaccuracies.

Furthermore, the interpretation of model-derived parameters should be approached with caution. Although the model demonstrated good spectral fits across groups, some parameter estimates exhibited high standard deviation during the fitting process, indicating low parameter identifiability and suggesting the possibility that different combinations of parameters could yield similarly accurate fits to the data. This issue, common in complex biophysical modelling ^85^, highlights the need for caution when drawing mechanistic conclusions from individual parameter values. On the other hand, the low correlation of CRS-R index and the model parameter values and signal complexity measures indicates a key distinction between the neural information captured by the model and behavioral scales. The behavioral scoring might be too nuanced and influenced by too many factors (e.g. motor and sensory deficits, patient fluctuations) for a linear correlation with a specific neural parameter to emerge. The effect of TSO further illustrates this dissociation: while there is no effect on the model parameters themselves, it did influence dynamic output measures such as the signal complexity. Consequently, while the model robustly differentiates diagnostic categories, linking its parameters more closely to bedside clinic assessment remains a challenge requiring further investigation.

The model’s ability to preserve key features of signal complexity in its outputs could be a motivating factor for extending its application to simulating brain stimulation and pharmacological interventions in future studies. As has been stated in the literature, any kind of perturbational analysis on a neural field model is only as valid as the model’s ability to capture relevant features of the brain dynamics ^86^. Therefore, the strong capacity of the baseline biophysical model to capture relevant aspects of whole-brain dynamics makes the simulation of external interventions more feasible and potentially reliable. In neural field models, neural stimulation effects could be implemented as a controlled external input to specific neural populations, described by the waveform and time course of induced electric fields, combined with population-based neural dynamics, and plasticity rules ^65,87^. The individualized virtual clinical trials generated by the models would enable robust simulation of post-simulation effects on EEG activity and thalamocortical loop parameters. Pharmacological interventions could also potentially be simulated in patient models using approaches similar to those used by Alnagger *et al.* and Mindlin et al. ^86,88^ on fMRI data, where parameter changes between placebo and drug conditions were used as modulating factors in the model to simulate treatment.

## Conclusion

We demonstrated the feasibility of using neural field models to fit high-density EEG recordings from patients with DoC, and their ability to shed light on the specific thalamocortical circuits that contribute to aberrant power spectra. Moreover, we showed the remarkable capacity of the model to generate synthetic signals that closely mimic complexity features of the empirical recordings. The model’s ability to detect impaired communication between neuronal populations, combined with its capacity to simulate EEG time series, makes it a strong candidate for providing mechanistic insight into individual patient impairments, estimating proximity to criticality, and testing personalized treatment strategies aimed at restoring consciousness.

## Author Contributions

Conceptualization: LC, PN, JS, JA

Methodology: LC, PN, PT

Software: LC, PN, PT

Formal analysis: LC

Investigation: LC, AT, OG

Resources: OG, JS

Data Curation and Preprocessing: JA, LB

Writing—original draft: LC, PN

Writing—review & editing: PT, NLNA, IM, GJML, LB, SL, LN, BR, AT, JA, OG, JS

Visualization: LC, PN

Supervision: PN, JS

## Acknowledgments

We thank the patients, their families and the control subjects for participating in our study. The authors extend their gratitude to the entire staff of the ICU and Nuclear Medicine and Radiodiagnostic departments at the University Hospital of Liège. We are especially thankful to the members of the Coma Science Group for their assistance with the clinical evaluations.

This work was supported by the University and University Hospital of Liège, the Belgian National Funds for Scientific Research (FRS-FNRS), the FNRS PDR project (T.0134.21), the FNRS MIS project (F.4521.23), the FLAG-ERA JTC2021 project ModelDXConsciousness (Human Brain Project Partnering Project), the program Investissements d’avenir ANR-10-IAIHU-06, JTC the fund Generet, the King Baudouin Foundation, the BIAL Foundation, the Mind Science Foundation, the European Commission, the Fondation Leon Fredericq, the Mind-Care foundation, the National Natural Science Foundation of China (Joint Research Project 81471100), the European Foundation of Biomedical Research FERB Onlus, and the Horizon 2020 MSCA – Research and Innovation Staff Exchange DoC-Box project (HORIZON-MSCA-2022-SE-01-01; 101131344). NA is a research fellow, OG and AT are research associates, and SL is research director at the F.R.S.-FNRS. JA is postdoctoral fellow at the FWO (1265522N).

## References

1. Zasler, N. D., Aloisi, M., Contrada, M. & Formisano, R. Disorders of consciousness terminology: history, evolution and future directions. Brain Inj. 33, 1684–1689 (2019).

2. Posner, J. B., Saper, C. B., Schiff, N. & Plum, F. Plum and Posner’s Diagnosis of Stupor and Coma. (Oxford University Press, 2008). doi:10.1093/med/9780195321319.001.0001

3. Giacino, J. T. et al. The minimally conscious state. Neurology 58, 349–353 (2002).

4. Laureys, S. et al. Unresponsive wakefulness syndrome: a new name for the vegetative state or apallic syndrome. BMC Med. 8, 68 (2010).

5. Whyte, C. J., Redinbaugh, M. J., Shine, J. M. & Saalmann, Y. B. Thalamic contributions to the state and contents of consciousness. Neuron 112, 1611–1625 (2024).

6. Redinbaugh, M. J. et al. Thalamus Modulates Consciousness via Layer-Specific Control of Cortex. Neuron 106, 66–75.e12 (2020).

7. Schiff, N. D. Central thalamic contributions to arousal regulation and neurological disorders of consciousness. Ann. N. Y. Acad. Sci. 1129, 105–118 (2008).

8. Schiff, N. D. Recovery of consciousness after brain injury: a mesocircuit hypothesis. Trends Neurosci. 33, 1–9 (2010).

9. Forgacs, P. B. et al. Dynamic regimes of neocortical activity linked to corticothalamic integrity correlate with outcomes in acute anoxic brain injury after cardiac arrest. Ann. Clin. Transl. Neurol. 4, 119–129 (2017).

10. Tasserie, J. et al. Deep brain stimulation of the thalamus restores signatures of consciousness in a nonhuman primate model. Sci. Adv. 8, 1–17 (2022).

11. Schiff, N. D. et al. Behavioural improvements with thalamic stimulation after severe traumatic brain injury. Nature 448, 600–603 (2007).

12. Dutta, R. R. et al. Neuromodulation and Disorders of Consciousness: Systematic Review and Pathophysiology. Neuromodulation 28, 380–400 (2025).

13. Gorka, S. M. et al. Alterations in large-scale resting-state network nodes following transcranial focused ultrasound of deep brain structures. Front. Hum. Neurosci. 18, 1–11 (2024).

14. Cain, J. A. et al. Ultrasonic thalamic stimulation in chronic disorders of consciousness. Brain Stimul. 14, 301–303 (2021).

15. Monti, M. M., Schnakers, C., Korb, A. S., Bystritsky, A. & Vespa, P. M. Non-Invasive Ultrasonic Thalamic Stimulation in Disorders of Consciousness after Severe Brain Injury: A First-in-Man Report. Brain Stimul. 9, 940–941 (2016).

16. Whyte, J. et al. Predictors of outcome in prolonged posttraumatic disorders of consciousness and assessment of medication effects: A multicenter study. Arch. Phys. Med. Rehabil. 86, 453–462 (2005).

17. Giacino, J. T., Fins, J. J., Laureys, S. & Schiff, N. D. Disorders of consciousness after acquired brain injury: the state of the science. Nat. Rev. Neurol. 10, 99–114 (2014).

18. Kalmar, K. & Giacino, J. The JFK coma recovery scale—revised. Neuropsychol. Rehabil. 15, 454–460 (2005).

19. Kondziella, D. et al. European Academy of Neurology guideline on the diagnosis of coma and other disorders of consciousness. Eur. J. Neurol. 27, 741–756 (2020).

20. Sitt, J. D. et al. Large scale screening of neural signatures of consciousness in patients in a vegetative or minimally conscious state. Brain 137, 2258–2270 (2014).

21. Annen, J. et al. Cerebral electrometabolic coupling in disordered and normal states of consciousness. Cell Rep. 42, 112854 (2023).

22. Engemann, D. A. et al. Robust EEG-based cross-site and cross-protocol classification of states of consciousness. Brain 141, 3179–3192 (2018).

23. Comanducci, A. et al. Clinical and advanced neurophysiology in the prognostic and diagnostic evaluation of disorders of consciousness: review of an IFCN-endorsed expert group. Clin. Neurophysiol. 131, 2736–2765 (2020).

24. Cook, B. J., Peterson, A. D. H., Woldman, W. & Terry, J. R. Neural Field Models: A mathematical overview and unifying framework. Math. Neurosci. Appl. Volume 2, 1–67 (2022).

25. Robinson, P. A., Loxley, P. N., O’Connor, S. C. & Rennie, C. J. Modal analysis of corticothalamic dynamics, electroencephalographic spectra, and evoked potentials. Phys. Rev. E 63, 041909 (2001).

26. Robinson, P. A., Rennie, C. J. & Rowe, D. L. Dynamics of large-scale brain activity in normal arousal states and epileptic seizures. *Phys*. Rev. E 65, 041924 (2002).

27. Robinson, P. A., Rennie, C. J., Rowe, D. L. & O’Connor, S. C. Estimation of multiscale neurophysiologic parameters by electroencephalographic means. Hum. Brain Mapp. 23, 53–72 (2004).

28. Abeysuriya, R. G., Rennie, C. J. & Robinson, P. A. Physiologically based arousal state estimation and dynamics. J. Neurosci. Methods 253, 55–69 (2015).

29. Rennie, C. J., Robinson, P. A. & Wright, J. J. Unified neurophysical model of EEG spectra and evoked potentials. Biol. Cybern. 86, 457–471 (2002).

30. Abeysuriya, R. G., Rennie, C. J. & Robinson, P. A. Prediction and verification of nonlinear sleep spindle harmonic oscillations. J. Theor. Biol. 344, 70–77 (2014).

31. Breakspear, M. et al. A unifying explanation of primary generalized seizures through nonlinear brain modeling and bifurcation analysis. Cereb. Cortex 16, 1296–1313 (2006).

32. van Albada, S. J., Kerr, C. C., Chiang, A. K. I., Rennie, C. J. & Robinson, P. A. Neurophysiological changes with age probed by inverse modeling of EEG spectra. Clin. Neurophysiol. 121, 21–38 (2010).

33. Assadzadeh, S. et al. Method for quantifying arousal and consciousness in healthy states and severe brain injury via EEG-based measures of corticothalamic physiology. J. Neurosci. Methods 398, 109958 (2023).

34. Sarasso, S. et al. Consciousness and complexity: a consilience of evidence. Neurosci. Conscious. 2021, 1–24 (2021).

35. Schartner, M. et al. Complexity of multi-dimensional spontaneous EEG decreases during propofol induced general anaesthesia. PLoS One 10, (2015).

36. Medel, V., Irani, M., Crossley, N., Ossandón, T. & Boncompte, G. Complexity and 1/f slope jointly reflect brain states. Sci. Rep. 13, 21700 (2023).

37. Casali, A. G. et al. A Theoretically Based Index of Consciousness Independent of Sensory Processing and Behavior. Sci. Transl. Med. 5, (2013).

38. Bandt, C. & Pompe, B. Permutation Entropy: A Natural Complexity Measure for Time Series. Phys. Rev. Lett. 88, 4 (2002).

39. Inouye, T. et al. Quantification of EEG irregularity by use of the entropy of the power spectrum. Electroencephalogr. Clin. Neurophysiol. 79, 204–210 (1991).

40. King, J.-R. et al. Information Sharing in the Brain Indexes Consciousness in Noncommunicative Patients. Curr. Biol. 23, 1914–1919 (2013).

41. Sitt, J. D. et al. Large scale screening of neural signatures of consciousness in patients in a vegetative or minimally conscious state. Brain 137, 2258–2270 (2014).

42. Annen, J. et al. Diagnostic accuracy of the CRS-R index in patients with disorders of consciousness. Brain Inj. 33, 1409–1412 (2019).

43. Mashour, G. A., Roelfsema, P., Changeux, J. P. & Dehaene, S. Conscious Processing and the Global Neuronal Workspace Hypothesis. Neuron 105, 776–798 (2020).

44. Dehaene, S. & Naccache, L. Towards a cognitive neuroscience of consciousness: Basic evidence and a workspace framework. Cognition 79, 1–37 (2001).

45. Tononi, G., Boly, M., Massimini, M. & Koch, C. Integrated information theory: From consciousness to its physical substrate. Nat. Rev. Neurosci. 17, 450–461 (2016).

46. Koch, C., Massimini, M., Boly, M. & Tononi, G. Neural correlates of consciousness: Progress and problems. Nat. Rev. Neurosci. 17, 307–321 (2016).

47. Cogitate Consortium et al. Adversarial testing of global neuronal workspace and integrated information theories of consciousness. Nature 642, (2025).

48. Melloni, L. et al. An adversarial collaboration protocol for testing contrasting predictions of global neuronal workspace and integrated information theory. PLoS One 18, 1–28 (2023).

49. Mudrik, L. et al. Unpacking the complexities of consciousness: Theories and reflections. Neurosci. Biobehav. Rev. 170, 106053 (2025).

50. Albantakis, L., et al. Integrated information theory (IIT) 4.0: Formulating the properties of phenomenal existence in physical terms. PLoS Computational Biology 19, (2023).

51. Wannez, S., Heine, L., Thonnard, M., Gosseries, O. & Laureys, S. The repetition of behavioral assessments in diagnosis of disorders of consciousness. Ann. Neurol. 81, 883–889 (2017).

52. Chennu, S. et al. Brain networks predict metabolism, diagnosis and prognosis at the bedside in disorders of consciousness. Brain 140, 2120–2132 (2017).

53. Bekinschtein, T. A. et al. Neural signature of the conscious processing of auditory regularities. Proc. Natl. Acad. Sci. U. S. A. 106, 1672–1677 (2009).

54. Thibaut, A. et al. Preservation of Brain Activity in Unresponsive Patients Identifies MCS Star. Ann. Neurol. 90, 89–100 (2021).

55. Manasova, D. et al. Dynamics of EEG Microstates Change Across the Spectrum of Disorders of Consciousness. Brain Topogr. 38, 1–12 (2025).

56. Manasova, D. et al. Integrative Electrophysiology and Neuroimaging Approach in Assessing Disorders of Consciousness: A Multimodal Multicentric Machine Learning Study. (2024).

57. Rohaut, B. et al. Multimodal assessment improves neuroprognosis performance in clinically unresponsive critical-care patients with brain injury. Nat. Med. 30, 2349–2355 (2024).

58. Engemann, D. et al. Automated Measurement and Prediction of Consciousness in Vegetative and Minimally Conscious Patients. ICML Work. Stat. Mach. Learn. Neurosci. (Stamlins 2015) (2015).

59. Robinson, P. A. et al. Prediction of electroencephalographic spectra from neurophysiology. Phys. Rev. E 63, 021903 (2001).

60. Srinivasan, R., Nunez, P. L. & Silberstein, R. B. Spatial filtering and neocortical dynamics: estimates of EEG coherence. IEEE Trans. Biomed. Eng. 45, 814–826 (1998).

61. Van Boxtel, A. Optimal signal bandwidth for the recording of surface EMG activity of facial, jaw, oral, and neck muscles. Psychophysiology 38, S004857720199016X (2001).

62. Goncharova, I.., McFarland, D.., Vaughan, T.. & Wolpaw, J.. EMG contamination of EEG: spectral and topographical characteristics. Clin. Neurophysiol. 114, 1580–1593 (2003).

63. Rowe, D. L., Robinson, P. A. & Rennie, C. J. Estimation of neurophysiological parameters from the waking EEG using a biophysical model of brain dynamics. J. Theor. Biol. 231, 413–433 (2004).

64. Abeysuriya, R. G. & Robinson, P. A. Real-time automated EEG tracking of brain states using neural field theory. J. Neurosci. Methods 258, 28–45 (2016).

65. Sanz-Leon, P. et al. NFTsim: Theory and Simulation of Multiscale Neural Field Dynamics. PLOS Comput. Biol. 14, e1006387 (2018).

66. Abásolo, D., Hornero, R., Gómez, C., García, M. & López, M. Analysis of EEG background activity in Alzheimer’s disease patients with Lempel-Ziv complexity and central tendency measure. Med. Eng. Phys. 28, 315–322 (2006).

67. Benjamini, Y. & Hochberg, Y. Controlling the false discovery rate: a practical and powerful approach to multiple testing. J. R. Stat. Soc. 57, 289–300 (1995).

68. Edlow, B. L., Claassen, J., Schiff, N. D. & Greer, D. M. Recovery from disorders of consciousness: mechanisms, prognosis and emerging therapies. Nat. Rev. Neurol. 17, 135–156 (2021).

69. Lutkenhoff, E. S. et al. Thalamic and extrathalamic mechanisms of consciousness after severe brain injury. Ann. Neurol. 78, 68–76 (2015).

70. Annen, J. et al. Function–structure connectivity in patients with severe brain injury as measured by MRI-DWI and FDG-PET. Hum. Brain Mapp. 37, 3707–3720 (2016).

71. Chen, P. et al. Abnormal Effective Connectivity of the Anterior Forebrain Regions in Disorders of Consciousness. Neurosci. Bull. 34, 647–658 (2018).

72. Tewarie, P. K. B., Tjepkema-Cloostermans, M. C., Abeysuriya, R. G., Hofmeijer, J. & van Putten, M. J. A. M. Preservation of thalamocortical circuitry is essential for good recovery after cardiac arrest. PNAS Nexus 2, 1–8 (2023).

73. Magrassi, L. et al. Single unit activities recorded in the thalamus and the overlying parietal cortex of subjects affected by disorders of consciousness. PLoS One 13, e0205967 (2018).

74. Fernández-Espejo, D. et al. Diffusion weighted imaging distinguishes the vegetative state from the minimally conscious state. Neuroimage 54, 103–112 (2011).

75. Cui, Y. et al. Subdivisions of the posteromedial cortex in disorders of consciousness. NeuroImage Clin. 20, 260–266 (2018).

76. Panda, R. et al. Whole-brain analyses indicate the impairment of posterior integration and thalamo-frontotemporal broadcasting in disorders of consciousness. Hum. Brain Mapp. 44, 4352–4371 (2023).

77. Wu, G. R. et al. Modulation of the spontaneous hemodynamic response function across levels of consciousness. Neuroimage 200, 450–459 (2019).

78. Stephan, B. C. M. et al. The neuropathological profile of mild cognitive impairment (MCI): a systematic review. Mol. Psychiatry 17, 1056–1076 (2012).

79. Naro, A. et al. Shedding new light on disorders of consciousness diagnosis: The dynamic functional connectivity. Cortex 103, 316–328 (2018).

80. Cocchi, L., Gollo, L. L., Zalesky, A. & Breakspear, M. Criticality in the brain: A synthesis of neurobiology, models and cognition. Prog. Neurobiol. 158, 132–152 (2017).

81. Luppi, A. I. et al. Computational modelling in disorders of consciousness: Closing the gap towards personalised models for restoring consciousness. Neuroimage 275, (2023).

82. Tagliazucchi, E. et al. Large-scale signatures of unconsciousness are consistent with a departure from critical dynamics. J. R. Soc. Interface 13, (2016).

83. Koch, C., Massimini, M., Boly, M. & Tononi, G. Posterior and anterior cortex — where is the difference that makes the difference? Nat. Rev. Neurosci. 17, 666–666 (2016).

84. Kayser, J. & Tenke, C. E. On the benefits of using surface Laplacian (current source density) methodology in electrophysiology. Int. J. Psychophysiol. 97, 171–173 (2015).

85. Gutenkunst, R. N. et al. Universally Sloppy Parameter Sensitivities in Systems Biology Models. PLoS Comput. Biol. 3, e189 (2007).

86. Mindlin, I. et al. Whole brain modelling for simulating pharmacological interventions on patients with disorders of consciousness. *Commun*. Biol. 7, 1176 (2024).

87. Fung, P. K. & Robinson, P. A. Neural field theory of synaptic metaplasticity with applications to theta burst stimulation. J. Theor. Biol. 340, 164–176 (2014).

88. Alnagger, N. L. N. et al. A virtual clinical trial of psychedelics to treat patients with disorders of consciousness. (2024). doi:10.1101/2024.08.16.608251

