## Supplementary Material for "Individualised electrophysiological neural field models for the assessment of thalamocortical function in disorders of consciousness: a multicentre study"

**Figure S1. Comparison of the parameters fitted to the power spectrum for the three subject groups across channels around different regions in the Liège dataset.** This figure shows the parameters derived from the corticothalamic model for three groups: healthy controls (HC), minimally conscious state (MCS), and unresponsive wakefulness syndrome (UWS) across frontal, parietal, and whole-brain EEG channels. A consistent pattern is observed across all regions, with significant differences between healthy controls and patients in the following model parameters: 1) $G_{ese}$, the gain of the excitatory corticothalamic loop was lower in DoC patients; 2) $G_{esre}$, the gain of the inhibitory corticothalamic loop is higher in DoC patients; 3) $G_{srs}$, the gain of the intrathalamic loop was higher in DoC patients; 4) $t_{0}$, the delay between cortical and thalamic regions was longer in DOC patients; 5) the EMG component amplitude was larger in DoC patients.

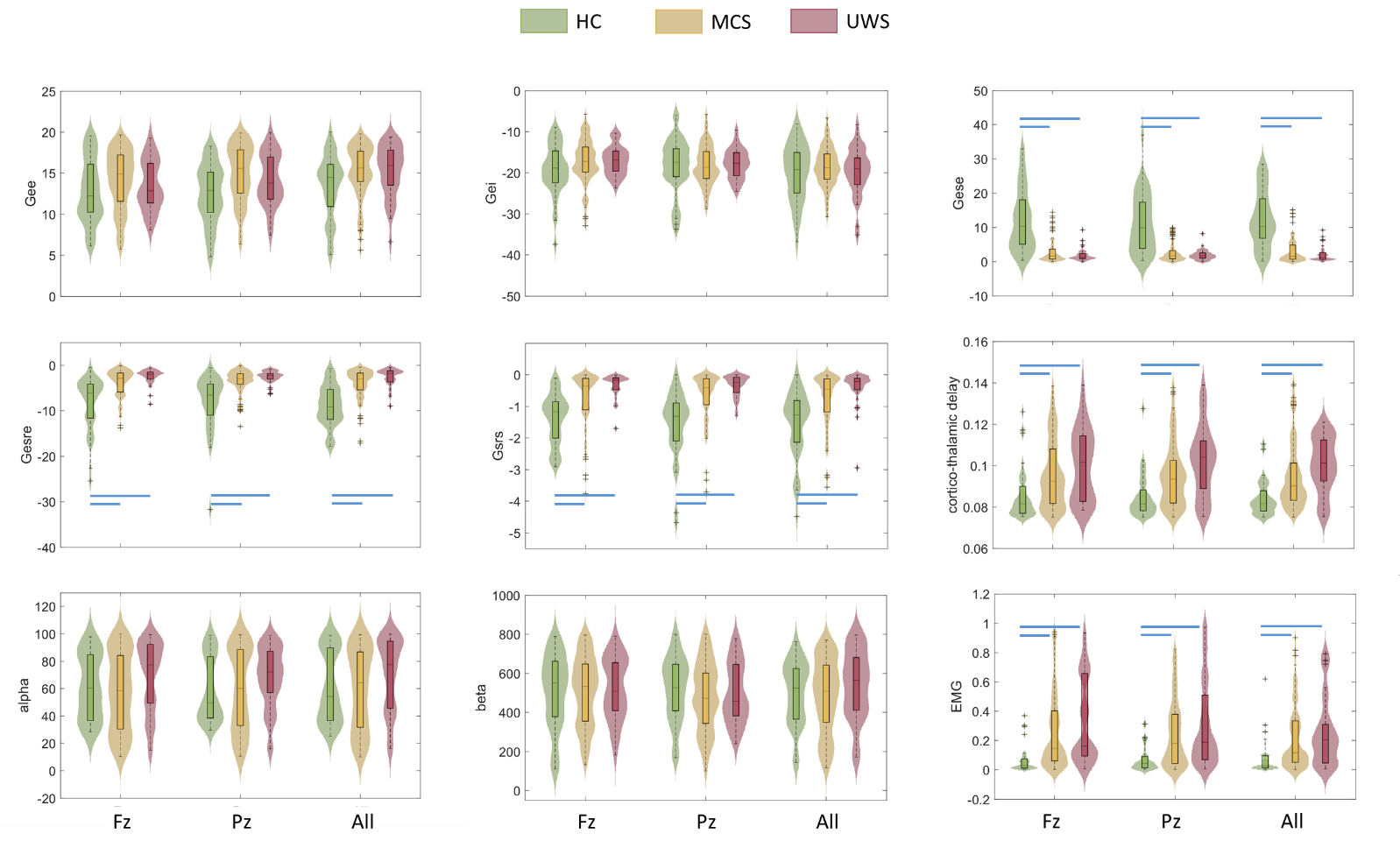

**Figure S2. Comparison for the parameters fitted to the power spectrum for the three subject groups across channels of different regions in the Paris dataset.** Consistent with findings from the Liège dataset, significant group differences were observed in several model parameters when fitting for frontal, parietal and whole-brain recordings: 1) $G_{ese}$, the gain of the excitatory corticothalamic loop was lower in DoC patients; 2) $G_{esre}$, the gain of the inhibitory corticothalamic loop is higher in DoC patients; 3) $G_{srs}$, the gain of the intrathalamic loop was higher in DoC patients; 4) $t_{0}$, the delay between cortical and thalamic regions was longer in DoC patients; 5) the EMG component amplitude was larger in DoC patients. Additionally, $G_{ei}$, the gain between excitatory and inhibitory cortical populations, is higher in patients than in healthy controls in the frontal regions; the gain of the excitatory corticothalamic loop is lower in unresponsive wakefulness syndrome (UWS) than in the frontal region, parietal region, and whole-brain; the corticothalamic delay is lower in UWS than in minimally conscious state (MCS) in parietal and whole-brain recordings; the EMG component amplitude is larger in UWS than in MCS across all regions.

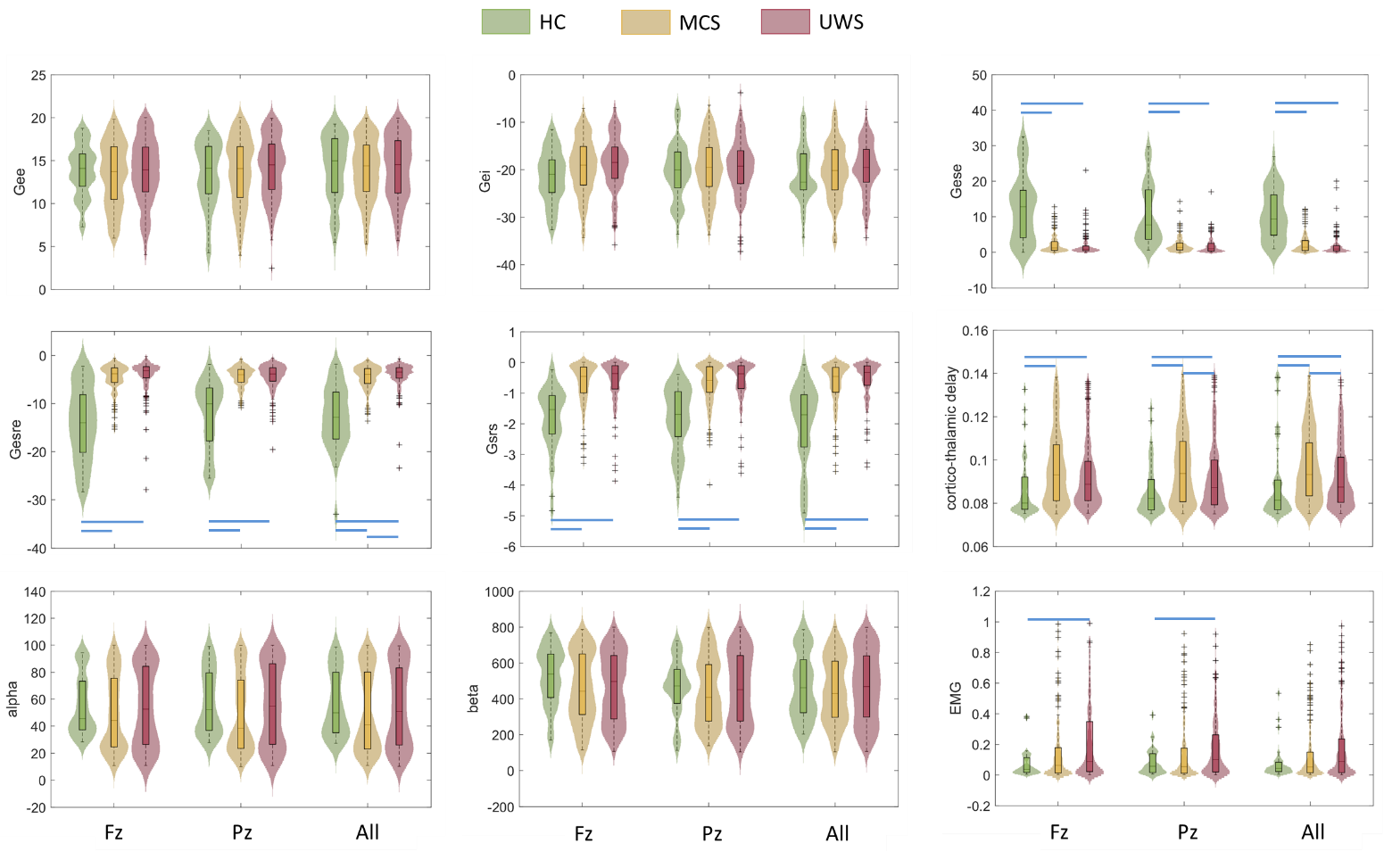

**Figure S3. Group comparisons of complexity measures in real recordings and simulations in the Liège dataset.** For sample entropy (SE) and permutation entropy (PE), signal complexity was higher in the healthy control group compared to the patient groups (MCS and UWS) in both real recordings and simulations, consistently across frontal and parietal channels. No significant differences were found between MCS and UWS groups for these two measures. For Lempel-Ziv complexity (LZC), no group differences were observed in the real recordings. However, in the simulations, LZC was higher in healthy controls compared to patient groups in both frontal and parietal channels. Additionally, for the parietal channels, LZC was significantly higher in the UWS group compared to the MCS group. HC: healthy control; MCS: minimally conscious state; UWS: unresponsive wakefulness syndrome.

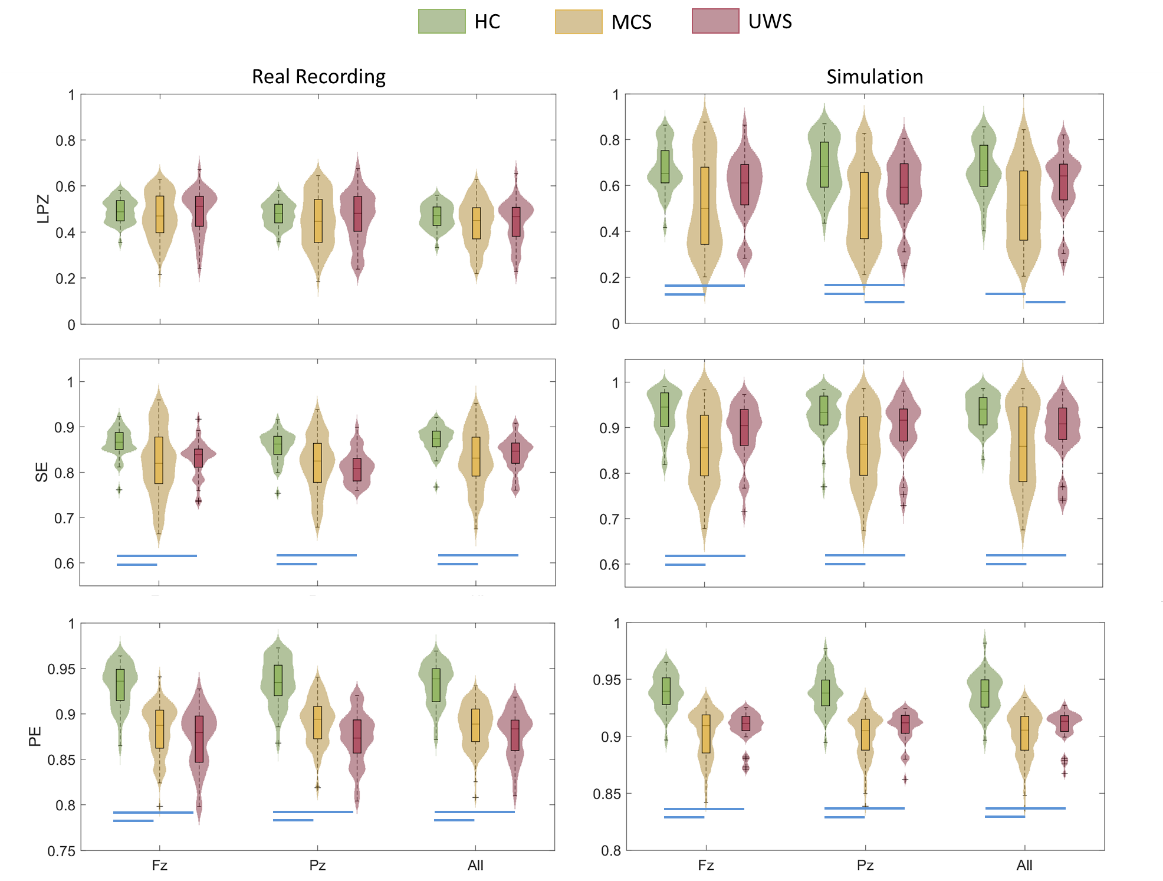

**Figure S4. Group comparison for complexity measures in real recordings and simulation of Paris dataset.** Across all three complexity measures and channel conditions, signal complexity was consistently higher in the healthy control group compared to the patient groups, in both real recordings and simulations. Additionally, PE was higher in the MCS group than in the UWS group for frontal and parietal channels in the real recordings, as well as for parietal channels in the simulations. HC: healthy control; MCS: minimally conscious state; UWS: unresponsive wakefulness syndrome. LZC: Lempel-Ziv complexity; SE: spectral entropy; PE: permutation entropy.

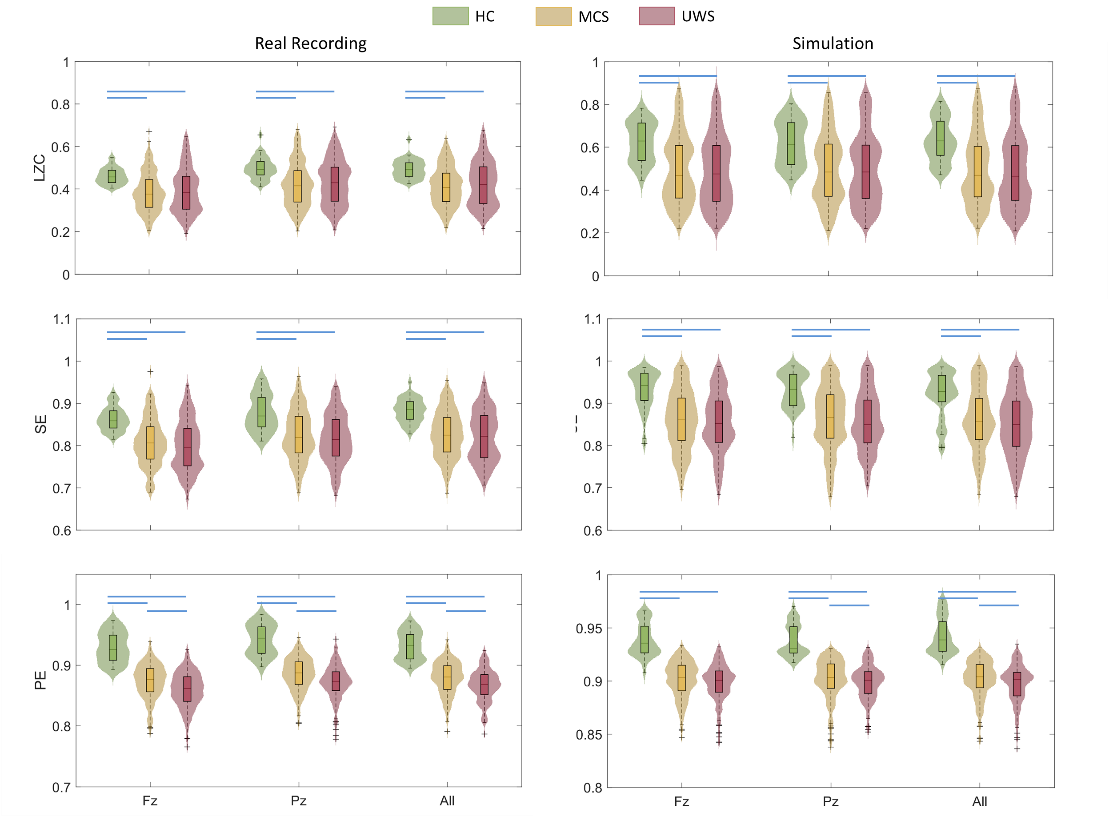

**Figure S5. Comparison of complexity measure values between the Time Since Onset (TSO) more than 1 year and the TSO less than 1 year groups in both datasets.** Blue lines indicate a significance level of *p*<0.05.

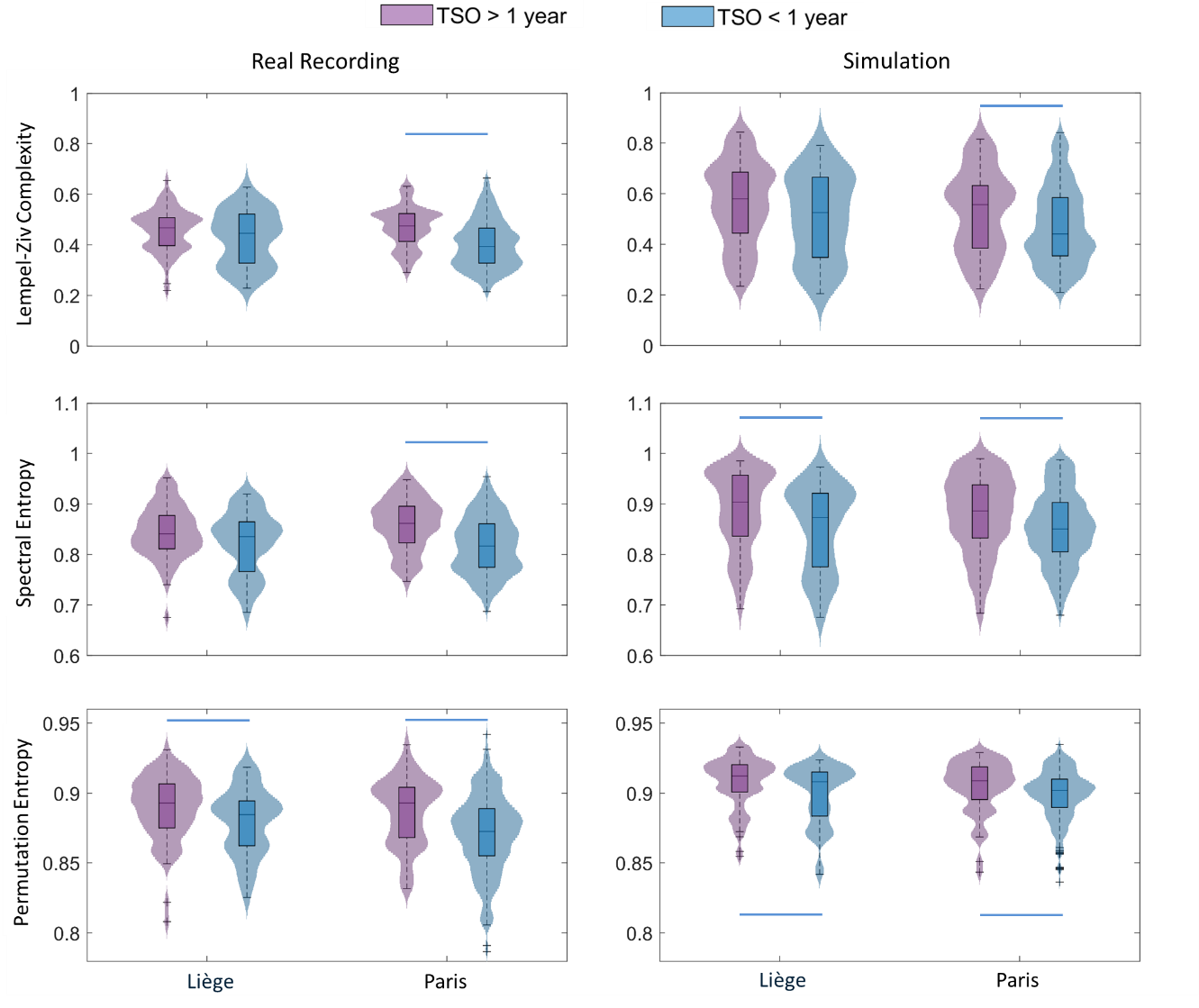

**Table S1. Statistical comparisons of group effects (HC, MCS, UWS) on model parameters across both datasets.** P-values were FDR-corrected for multiple comparisons (Kruskal-Wallis test). In addition to comparing parameter values across groups for the frontal and parietal channel models separately, we also computed the difference in parameter values between these two models for each subject, and tested group differences on these deltas across nine parameters. No significant group differences were found in any parameter, suggesting that model performance on frontal versus parietal channels did not differ significantly between subject groups. HC: healthy control; MCS: minimally conscious state; UWS: unresponsive wakefulness syndrome.

|  |  |  | $\boldsymbol{G}_{\boldsymbol{ee}}$ | $\boldsymbol{G}_{\boldsymbol{ei}}$ | $\boldsymbol{G}_{\boldsymbol{ese}}$ | $\boldsymbol{G}_{\boldsymbol{esre}}$ | $\boldsymbol{G}_{\boldsymbol{srs}}$ | $\boldsymbol{\alpha}$ | $\boldsymbol{\beta}$ | $\boldsymbol{t}_{\boldsymbol{0}}$ | $\boldsymbol{P}_{\boldsymbol{EMG}}$ |
| --- | --- | --- | --- | --- | --- | --- | --- | --- | --- | --- | --- |
| **Liège** | **Frontal** | $\boldsymbol{\chi}^{\boldsymbol{2}}$ | 3.519 | 3.586 | 44.274 | 36.901 | 34.199 | 5.889 | 0.436 | 17.622 | 33.243 |
|  |  | **p-value** | 0.194 | 0.194 | <0.001 | <0.001 | <0.001 | 0.079 | 0.804 | <0.001 | 0.004 |
|  | **Parietal** | $\boldsymbol{\chi}^{\boldsymbol{2}}$ | 11.295 | 0.308 | 38.845 | 34.707 | 38.677 | 2.726 | 1.869 | 24.066 | 26.105 |
|  |  | **p-value** | 0.005 | 0.857 | <0.001 | <0.001 | <0.001 | 0.329 | 0.442 | <0.001 | <0.001 |
|  | **Difference** | $\boldsymbol{\chi}^{\boldsymbol{2}}$ | 1.471 | 1.824 | 1.112 | 0.941 | 1.058 | 2.201 | 1.622 | 0.020 | 0.338 |
|  |  | **p-value** | 0.803 | 0.803 | 0.803 | 0.803 | 0.803 | 0.803 | 0.803 | 0.990 | 0.950 |
| **Paris** | **Frontal** | $\boldsymbol{\chi}^{\boldsymbol{2}}$ | 0.978 | 6.762 | 60.014 | 63.793 | 55.139 | 4.099 | 3.622 | 14.889 | 8.895 |
|  |  | **p-value** | 0.613 | 0.051 | <0.001 | <0.001 | <0.001 | 0.166 | 0.184 | 0.003 | 0.021 |
|  | **Parietal** | $\boldsymbol{\chi}^{\boldsymbol{2}}$ | 1.416 | 0.678 | 64.148 | 60.946 | 54.129 | 6.789 | 0.726 | 17.840 | 6.573 |
|  |  | **p-value** | 0.633 | 0.713 | <0.001 | <0.001 | <0.001 | 0.056 | 0.713 | <0.001 | 0.056 |
|  | **Difference** | $\boldsymbol{\chi}^{\boldsymbol{2}}$ | 0.258 | 0.728 | 1.882 | 0.576 | 5.791 | 0.219 | 0.804 | 1.609 | 0.164 |
|  |  | **p-value** | 0.921 | 0.921 | 0.921 | 0.921 | 0.498 | 0.921 | 0.921 | 0.921 | 0.921 |

**Table S2. Statistical comparisons of the pairwise interactions between groups (UWS, MCS, HC) of the model parameters in the two datasets (Mann-Whitney *U-*test). FDR corrected for the number of comparisons.** HC: healthy control; MCS: minimally conscious state; UWS: unresponsive wakefulness syndrome.

|  |  |  |  | $\boldsymbol{G}_{\boldsymbol{ee}}$ | $\boldsymbol{G}_{\boldsymbol{ei}}$ | $\boldsymbol{G}_{\boldsymbol{ese}}$ | $\boldsymbol{G}_{\boldsymbol{esre}}$ | $\boldsymbol{G}_{\boldsymbol{srs}}$ | $\boldsymbol{\alpha}$ | $\boldsymbol{\beta}$ | $\boldsymbol{t}_{\boldsymbol{0}}$ | $\boldsymbol{P}_{\boldsymbol{EMG}}$ |
| --- | --- | --- | --- | --- | --- | --- | --- | --- | --- | --- | --- | --- |
| **Liège** | **Frontal** | **HC-MCS** | ***U-*value** | 2156 | 1661 | 3361 | 1026 | 1284 | 2259 | 2428 | 1728 | 1472 |
|  |  |  | ***p*-value** | 0.532 | 0.001 | <0.001 | <0.001 | <0.001 | 0.834 | 0.677 | 0.003 | <0.001 |
|  |  | **HC-UWS** | ***U-*value** | 1243 | 1171 | 1790 | 832 | 817 | 1184 | 1279 | 991 | 868 |
|  |  |  | ***p*-value** | 0.462 | 0.142 | <0.001 | <0.001 | <0.001 | 0.166 | 0.689 | <0.001 | <0.001 |
|  |  | **MCS-UWS** | ***U-*value** | 5341 | 5172 | 5257 | 4830 | 4795 | 4758 | 5049 | 4950 | 4881 |
|  |  |  | ***p*-value** | 0.426 | 0.946 | 0.638 | 0.151 | 0.137 | 0.137 | 0.638 | 0.385 | 0.221 |
|  | **Parietal** | **HC-MCS** | ***U-*value** | 2117 | 1838 | 3448 | 984 | 1229 | 2218 | 2245 | 1753 | 1532 |
|  |  |  | ***p*-value** | 0.408 | 0.022 | <0.001 | <0.001 | <0.001 | 0.718 | 0.742 | 0.006 | <0.001 |
|  |  | **HC-UWS** | ***U-*value** | 1137 | 1341 | 1749 | 870 | 819 | 1167 | 1359 | 949 | 921 |
|  |  |  | ***p-*value** | 0.057 | 0.751 | <0.001 | <0.001 | <0.001 | 0.110 | 0.670 | <0.001 | <0.001 |
|  |  | **MCS-UWS** | ***U-*value** | 5339 | 5087 | 5231 | 4907 | 4783 | 4926 | 5047 | 4824 | 4952 |
|  |  |  | ***p-*value** | 0.434 | 0.676 | 0.676 | 0.373 | 0.209 | 0.373 | 0.648 | 0.209 | 0.393 |
| **Paris** | **Frontal** | **HC-MCS** | ***U-*value** | 4291 | 3414 | 6512 | 1585 | 1692 | 4818 | 4788 | 2805 | 3723 |
|  |  |  | ***p-*value** | 0.605 | 0.071 | <0.001 | <0.001 | <0.001 | 0.071 | 0.071 | <0.001 | 0.315 |
|  |  | **HC-UWS** | ***U-*value** | 3833 | 2981 | 6226 | 1379 | 1483 | 4105 | 4405 | 2800 | 3008 |
|  |  |  | ***p-*value** | 0.965 | 0.016 | <0.001 | <0.001 | <0.001 | 0.492 | 0.118 | 0.003 | 0.017 |
|  |  | **MCS-UWS** | ***U-*value** | 31742 | 31677 | 35297 | 30224 | 31329 | 31400 | 32486 | 34075 | 30445 |
|  |  |  | ***p-*value** | 0.383 | 0.383 | 0.051 | 0.051 | 0.286 | 0.286 | 0.857 | 0.286 | 0.064 |
|  | **Parietal** | **HC-MCS** | ***U-*value** | 3982 | 3860 | 6712 | 1466 | 1787 | 4944 | 4398 | 2765 | 4017 |
|  |  |  | ***p-*value** | 0.801 | 0.627 | <0.001 | <0.001 | <0.001 | 0.331 | 0.620 | <0.001 | 0.801 |
|  |  | **HC-UWS** | ***U-*value** | 3539 | 3565 | 6345 | 1425 | 1468 | 4060 | 3890 | 3124 | 3193 |
|  |  |  | ***p-*value** | 0.504 | 0.504 | <0.001 | <0.001 | <0.001 | 0.587 | 0.900 | 0.022 | 0.040 |
|  |  | **MCS-UWS** | ***U-*value** | 31711 | 32533 | 34546 | 31583 | 31103 | 30770 | 32052 | 35371 | 30517 |
|  |  |  | ***p-*value** | 0.417 | 0.895 | 0.112 | 0.395 | 0.190 | 0.112 | 0.594 | 0.044 | 0.112 |

**Table S3. Statistical comparison of model parameters between the Time Since Onset (TSO) of more than 1 year and the TSO of less than 1 year groups in both datasets.**

|  |  | $\boldsymbol{G}_{\boldsymbol{ee}}$ | $\boldsymbol{G}_{\boldsymbol{ei}}$ | $\boldsymbol{G}_{\boldsymbol{ese}}$ | $\boldsymbol{G}_{\boldsymbol{esre}}$ | $\boldsymbol{G}_{\boldsymbol{srs}}$ | $\boldsymbol{\alpha}$ | $\boldsymbol{\beta}$ | $\boldsymbol{t}_{\boldsymbol{0}}$ | $\boldsymbol{P}_{\boldsymbol{EMG}}$ |
| --- | --- | --- | --- | --- | --- | --- | --- | --- | --- | --- |
| **Liège** | ***U-*value** | 4059 | 3894 | 4299 | 3714 | 3899 | 4204 | 4124 | 3888 | 4052 |
|  | ***p-*value** | 0.643 | 0.643 | 0.428 | 0.487 | 0.643 | 0.487 | 0.643 | 0.643 | 0.643 |
| **Paris** | ***U-*value** | 10058 | 9704 | 10280 | 8709 | 9187 | 10939 | 11571 | 9217 | 10466 |
|  | ***p-*value** | 0.723 | 0.965 | 0.587 | 0.421 | 0.587 | 0.376 | 0.075 | 0.587 | 0.587 |

**Table S4. Correlation between model parameters and CRS-R index.**

|  |  | $\boldsymbol{G}_{\boldsymbol{ee}}$ | $\boldsymbol{G}_{\boldsymbol{ei}}$ | $\boldsymbol{G}_{\boldsymbol{ese}}$ | $\boldsymbol{G}_{\boldsymbol{esre}}$ | $\boldsymbol{G}_{\boldsymbol{srs}}$ | $\boldsymbol{\alpha}$ | $\boldsymbol{\beta}$ | $\boldsymbol{t}_{\boldsymbol{0}}$ | $\boldsymbol{P}_{\boldsymbol{EMG}}$ |
| --- | --- | --- | --- | --- | --- | --- | --- | --- | --- | --- |
| **Liège** | ***ρ*** | -0.044 | 0.120 | 0.237 | -0.254 | -0.094 | -0.068 | -0.042 | -0.081 | -0.063 |
|  | ***p-*value** | 0.658 | 0.617 | 0.051 | 0.051 | 0.651 | 0.651 | 0.658 | 0.651 | 0.651 |
| **Paris** | ***ρ*** | -0.057 | 0.058 | 0.009 | 0.024 | -0.053 | 0.008 | 0.043 | 0.027 | -0.016 |
|  | ***p-*value** | 0.291 | 0.285 | 0.870 | 0.659 | 0.325 | 0.883 | 0.423 | 0.613 | 0.767 |

**Table S5. Comparison of the complexity measures for frontal and parietal channels across groups (Kruskal-wallis test).** We computed the difference in complexity measures between the frontal and parietal channels and performed a Kruskal-Wallis test across the three subject groups. No significant differences were found for any of the three measures, indicating that signal complexity between frontal and parietal recordings did not differ across groups. HC: healthy control; MCS: minimally conscious state; UWS: unresponsive wakefulness syndrome. LZC: Lempel-Ziv complexity; SE: spectral entropy; PE: permutation entropy.

|  |  | **LZC** | | **SE** | | **PE** | |
| --- | --- | --- | --- | --- | --- | --- | --- |
|  |  | $\boldsymbol{\chi}^{\mathbf{2}}$ | ***p-*value** | $\boldsymbol{\chi}^{\mathbf{2}}$ | ***p-*value** | $\boldsymbol{\chi}^{\mathbf{2}}$ | ***p-*value** |
| **Liège** | **Frontal** | 2.254 | 0.324 | 17.483 | <0.001 | 58.611 | <0.001 |
|  | **Parietal** | 2.203 | 0.332 | 19.212 | <0.001 | 61.268 | <0.001 |
|  | **Difference** | 0.956 | 0.620 | 4.587 | 0.101 | 4.58 | 0.101 |
| **Paris** | **Frontal** | 31.5 | <0.001 | 40.832 | <0.001 | 102.496 | <0.001 |
|  | **Parietal** | 23.9 | <0.001 | 31.882 | <0.001 | 102.197 | <0.001 |
|  | **Difference** | 0.902 | 0.637 | 0.598 | 0.742 | 0.729 | 0.695 |

**Table S6. Pairwise comparison for complexity measures in frontal and parietal channels for real recordings and simulation of two datasets (Mann-Whitney *U-*test)**. HC: healthy control; MCS: minimally conscious state; UWS: unresponsive wakefulness syndrome. LZC: Lempel-Ziv complexity; SE: spectral entropy; PE: permutation entropy.

|  |  |  |  | **LZC** | | **SE** | | **PE** | |
| --- | --- | --- | --- | --- | --- | --- | --- | --- | --- |
|  |  |  |  | ***U-*value** | ***p-*value** | ***U-*value** | ***p-*value** | ***U-*value** | ***p-*value** |
| **Liège** | **Recordings** | **Frontal** | **HC-MCS** | 2526 | 0.202 | 2989 | <0.001 | 3585 | <0.001 |
|  |  |  | **HC-UWS** | 1264 | 0.564 | 1654 | <0.001 | 1836 | <0.001 |
|  |  |  | **MCS-UWS** | 4975 | 0.273 | 5004 | 0.356 | 1833 | 0.193 |
|  |  | **Parietal** | **HC-MCS** | 2554 | 0.152 | 2944 | <0.001 | 3517 | <0.001 |
|  |  |  | **HC-UWS** | 1362 | 0.860 | 1707 | <0.001 | 1862 | <0.001 |
|  |  |  | **MCS-UWS** | 5064 | 0.344 | 5327 | 0.323 | 1543 | 0.003 |
|  | **Simulation** | **Frontal** | **HC-MCS** | 3042 | <0.001 | 3170 | <0.001 | 3698 | <0.001 |
|  |  |  | **HC-UWS** | 1518 | 0.016 | 1573 | 0.002 | 1857 | <0.001 |
|  |  |  | **MCS-UWS** | 4821 | 0.045 | 4804 | 0.035 | 4870 | 0.086 |
|  |  | **Parietal** | **HC-MCS** | 3198 | <0.001 | 3161 | <0.001 | 3656 | <0.001 |
|  |  |  | **HC-UWS** | 1546 | 0.006 | 1509 | 0.022 | 1842 | <0.001 |
|  |  |  | **MCS-UWS** | 4751 | 0.015 | 4699 | 0.006 | 4868 | 0.084 |
| **Paris** | **Recordings** | **Frontal** | **HC-MCS**  **HC-UWS** | 6151 | <0.001 | 6255 | <0.001 | 7110 | <0.001 |
|  |  |  |  | 5474 | <0.001 | 5811 | <0.001 | 6854 | <0.001 |
|  |  |  | **MCS-UWS** | 32948 | 0.765 | 28937 | 0.198 | 25775 | <0.001 |
|  |  | **Parietal** | **HC-MCS** | 5875 | <0.001 | 5897 | <0.001 | 7110 | <0.001 |
|  |  |  | **HC-UWS**  **MCS-UWS** | 5259 | <0.001 | 5685 | <0.001 | 6844 | <0.001 |
|  |  |  |  | 32213 | 0.643 | 29138 | 0.281 | 25772 | <0.001 |
|  | **Simulation** | **Frontal** | **HC-MCS**  **HC-UWS** | 5948 | <0.001 | 6110 | <0.001 | 7268 | <0.001 |
|  |  |  |  | 5629 | <0.001 | 5834 | <0.001 | 6840 | <0.001 |
|  |  |  | **MCS-UWS** | 32862 | 0.834 | 29495 | 0.480 | 28413 | 0.067 |
|  |  | **Parietal** | **HC-MCS** | 5738 | <0.001 | 6021 | <0.001 | 7268 | <0.001 |
|  |  |  | **HC-UWS**  **MCS-UWS** | 5454 | <0.001 | 5819 | <0.001 | 6856 | <0.001 |
|  |  |  |  | 33075 | 0.667 | 29162 | 0.293 | 28015 | 0.025 |

**Table S7. Correlation analyses between empirical and simulated data (Spearman’s coefficient) for the frontal and parietal areas of the three complexity measures (LZC, SE and PE) in Liège and Paris datasets.** HC: healthy control; MCS: minimally conscious state; UWS: unresponsive wakefulness syndrome. LZC: Lempel-Ziv complexity; SE: spectral entropy; PE: permutation entropy.

|  |  |  | **LZC** | | **SE** | | **PE** | |
| --- | --- | --- | --- | --- | --- | --- | --- | --- |
|  |  |  | ***ρ*** | ***p-*value** | ***ρ*** | ***p-*value** | ***ρ*** | ***p-*value** |
| **Liège** | **Frontal** | **HC** | 0.669 | <0.001 | 0.594 | <0.001 | 0.903 | <0.001 |
|  |  | **MCS** | 0.625 | <0.001 | 0.700 | <0.001 | 0.820 | <0.001 |
|  |  | **UWS** | 0.261 | 0.063 | 0.492 | 0.004 | 0.485 | 0.004 |
|  | **Parietal** | **HC** | 0.762 | <0.001 | 0.680 | <0.001 | 0.825 | <0.001 |
|  |  | **MCS** | 0.550 | <0.001 | 0.694 | <0.001 | 0.758 | <0.001 |
|  |  | **UWS** | 0.343 | 0.016 | 0.576 | <0.001 | 0.389 | 0.025 |
| **Paris** | **Frontal** | **HC** | 0.644 | <0.001 | 0.764 | <0.001 | 0.805 | <0.001 |
|  |  | **MCS** | 0.484 | <0.001 | 0.602 | <0.001 | 0.682 | <0.001 |
|  |  | **UWS** | 0.313 | <0.001 | 0.466 | <0.001 | 0.598 | <0.001 |
|  | **Parietal** | **HC** | 0.715 | <0.001 | 0.740 | <0.001 | 0.712 | <0.001 |
|  |  | **MCS** | 0.394 | <0.001 | 0.544 | <0.001 | 0.693 | <0.001 |
|  |  | **UWS** | 0.277 | <0.001 | 0.385 | <0.001 | 0.544 | <0.001 |

**Table S8. Statistical comparison of complexity measures between the Time Since Onset (TSO) more than 1 year and the TSO less than 1 year groups in both datasets.** In Liège dataset, the SE value is higher in the TSO>1 group than in the TSO<1 group in simulation; and the PE value is higher in the TSO>1 group in both real recording and simulation. In Paris dataset, for all three measures in both real recording and model simulation, the signal complexity of the TSO>1 group is higher than the TSO<1 group. LZC: Lempel-Ziv complexity; SE: spectral entropy; PE: permutation entropy.

|  |  | **LZC** | | **SE** | | **PE** | |
| --- | --- | --- | --- | --- | --- | --- | --- |
|  |  | ***U-*value** | ***p-*value** | ***U-*value** | ***p-*value** | ***U-*value** | ***p-*value** |
| **Liège** | **Real recording** | 4222 | 0.132 | 4202 | 0.166 | 4342 | **0.025** |
|  | **Simulation** | 4239 | 0.107 | 4325 | **0.032** | 4319 | **0.035** |
| **Paris** | **Real recording** | 13238 | **<0.001** | 13167 | **<0.001** | 12480 | **<0.001** |
|  | **Simulation** | 11443 | **0.014** | 11512 | **0.011** | 11659 | **0.006** |

**Table S9. Correlation between the three complexity measures and CRS-R indexes in both datasets.** Significant correlations were found in PE value of the Paris dataset in both real recording and simulation. LZC: Lempel-Ziv complexity; SE: spectral entropy; PE: permutation entropy.

|  |  | **LZC** | | **SE** | | **PE** | |
| --- | --- | --- | --- | --- | --- | --- | --- |
|  |  | ***ρ*** | ***p-*value** | ***ρ*** | ***p-*value** | ***ρ*** | ***p-*value** |
| **Liège** | **Real recording** | -0.048 | 0.616 | -0.035 | 0.710 | 0.186 | 0.049 |
|  | **Simulation** | -0.062 | 0.514 | -0.004 | 0.969 | 0.011 | 0.912 |
| **Paris** | **Real recording** | 0.090 | 0.093 | 0.134 | **0.012** | 0.214 | **<0.001** |
|  | **Simulation** | 0.093 | 0.081 | 0.118 | **0.027** | 0.151 | **0.005** |
